# A novel dimeric FAP-targeting small molecule-radio conjugate with high and prolonged tumour uptake

**DOI:** 10.1101/2022.02.21.481260

**Authors:** Andrea Galbiati, Aureliano Zana, Matilde Bocci, Jacopo Millul, Abdullah Elsayed, Jacqueline Mock, Dario Neri, Samuele Cazzamalli

## Abstract

Imaging procedures based on small molecule-radio conjugates (SMRCs) targeting fibroblast activation protein (FAP) have recently emerged as a powerful tool for the diagnosis of a wide variety of tumours. However, the therapeutic potential of radiolabeled FAP-targeting agents is limited by their short residence time in neoplastic lesions. In this work, we present the development and *in vivo* characterization of BiOncoFAP, a new dimeric FAP-binding motif with extended tumour residence time and favorable tumour-to-organ ratio.

**Methods:** The binding properties of BiOncoFAP and its monovalent OncoFAP analogue were assayed against recombinant hFAP. Preclinical experiments with [^177^Lu]Lu-OncoFAP-DOTAGA (^177^Lu-OncoFAP) and [^177^Lu]Lu-BiOncoFAP-DOTAGA (^177^Lu-BiOncoFAP) were performed in mice bearing FAP-positive HT-1080 tumours.

**Results:** OncoFAP and BiOncoFAP displayed comparable sub-nanomolar dissociation constants towards hFAP in solution, but the bivalent BiOncoFAP bound more avidly to the target immobilized on solid supports. In a comparative biodistribution study, ^177^Lu-BiOncoFAP exhibited a more stable and prolonged tumour uptake than ^177^Lu-OncoFAP (∼20% ID/g vs ∼4% ID/g, at 24h p.i., respectively). Notably, ^177^Lu-BiOncoFAP showed favorable tumour-to-organ ratios with low kidney uptake. Both ^177^Lu-OncoFAP and ^177^Lu-BiOncoFAP displayed potent anti-tumour efficacy when administered at therapeutic doses in tumour bearing mice.

**Conclusions:** ^177^Lu-BiOncoFAP is a promising candidate for radioligand therapy of cancer, with favorable *in vivo* tumour-to-organ ratio, long tumour residence time and potent anti-cancer efficacy.

## Introduction

Small molecule-radio conjugates (SMRCs) are pharmaceutical products composed of a small organic ligand, acting as tumour targeting agent, and a radionuclide payload, that can be exploited both for diagnostic and therapeutic applications.(*1*–*3*) The “theranostic” potential of SMRCs, *id est* the possibility to perform imaging and therapy with the same product, facilitates the clinical development of this new class of drugs.(*4*–*7*) Patients who can predictably benefit from targeted radioligand therapy are accurately selected through dosimetry studies.(*8*) Lutathera®, a radioligand therapeutic targeting Somatostatin Receptor type 2 (SSTR-2), is the first SMRC product that gained marketing authorization for the therapy of neuroendocrine tumours (NETs).(*9*) The use of this drug has consistently shown high response rates and long median progression-free survival in a multicenter phase-III clinical trial.(*10*) More recently, a second product named ^177^Lu-PSMA-617 was shown to provide therapeutic benefit to PSMA-positive metastatic castration-resistant prostate cancer patients (mCRPC) in a large phase III clinical trial.(*11*) Radioligand therapy with ^177^Lu-PSMA-617 prolonged imaging-based progression-free survival and overall survival when added to standard care.(*11*)

In the last few years, a new category of pan-tumoural tumour-targeting SMRCs specific for Fibroblast Activation Protein (FAP) has been successfully implemented for the diagnosis of solid tumours.(*12*–*15*) FAP is a membrane-bound enzyme highly expressed on the surface of cancer-associated fibroblasts (CAFs) in the stroma of more than 90% of human epithelial cancers. FAP expression in healthy tissues is negligible.(*12,13,16,17*) We have recently reported the discovery of OncoFAP, the small molecule FAP-targeting agent with the highest affinity reported so far.(*18*) Proof-of-concept targeting studies with [^68^Ga]Ga-OncoFAP-DOTAGA (^68^Ga-OncoFAP), a PET tracer based on OncoFAP, have confirmed excellent biodistribution in patients with different primary and metastatic solid malignancies.(*19*)

Efficacy of radioligand therapeutics is strongly correlated to their residence time in tumours.(*9,20*–*23*) While Lutathera® and PSMA-617 are characterized by a sustained tumour residence time in patients (i.e., ∼61 hours for ^177^Lu-PSMA-617 and ∼88 hours for ^177^Lu-DOTATATE) (*24,25*), SMRCs based on FAP-targeting agents are typically cleared from solid lesions in few hours.(*26,27*) In preclinical biodistribution experiments, ^177^Lu-OncoFAP selectively localized on neoplastic lesions (∼38% ID/g, 1h after systemic administration), but half of the dose delivered to the tumour was lost within 8-12 hours.(*18*) A comparable tumour targeting performance and pharmacokinetic profile have been reported for other FAP-targeting SMRCs by Haberkorn and co-workers (e.g., the tumour uptake of [^177^Lu]Lu-FAPI-46 decreased from 12.5% ID/g at 1h to 2.5% ID/g at 24h after administration).(*28*) Importantly, a rapid washout from tumours is observed not only in mice but also in patients treated with [^177^Lu]Lu-FAPI-46.(*29*)

In an attempt to extend tumour residence time of FAP-targeting SMRCs and to maximize the exposure of cancer cells to biocidal radiation, we developed BiOncoFAP, a dimeric FAP-targeting OncoFAP-derivative. In this work, we describe the *in vitro* characterization of BiOncoFAP and we report the first preclinical biodistribution and therapy studies with a radiolabeled preparation of this novel dimeric FAP-targeting compound.

## Material and methods

Detailed experimental chemical and radio-chemical procedures are described in the **Supplemental material**, together with protocols of *in vitro* assays.

### In vitro Inhibition Assay on hFAP

Enzymatic activity of hFAP on the Z-Gly-Pro-AMC substrate was measured at room temperature on a microtiter plate reader, monitoring the fluorescence at an excitation wavelength of 360 nm and an emission wavelength of 465 nm. The reaction mixture contained substrate (20 μM), protein (200 pM, constant), assay buffer (50 mM Tris, 100 mM NaCl, and 1 mM EDTA, pH = 7.4), and inhibitors (compounds **1, 2, 4, 5**) with serial dilution from 1.67 μM to 800 fM, 1:2 in a total volume of 20 μL. Experiments were performed in triplicate, and the mean fluorescence values were fitted using Prism 7 (Y = Bottom + (Top Bottom)/(1 + ((X^HillSlope)/(IC50^HillSlope)))). The value is defined as the concentration of inhibitor required to reduce the enzyme activity by 50% after addition of the substrate.

### Affinity Measurement to hFAP by Fluorescence Polarization

Fluorescence polarization experiments were performed in 384-well plates (nonbinding, ps, f-bottom, black, high volume, 30 μL final volume). Stock solutions of proteins were serially diluted (1:2) with buffer (50 mM Tris, 100 mM NaCl, and 1 mM EDTA, pH = 7.4), while the final concentration of the binders (OncoFAP-Fluorescein and BiOncoFAP-Fluorescein) was kept constant at 10 nM. The fluorescence anisotropy was measured on a Tecan microtiter plate reader. Experiments were performed in triplicate, and the mean anisotropy values were fitted using Prism 7 ((Y = m1 + m2 × 0.5 × ((X + k + m3) sqrt((X + k + m3)^2 − 4 × X × k)), where k is the concentration of the fluorescent binder). Data are reported in the **Supplemental Material** [**Figure S2**].

### ELISA

Recombinant human FAP (1 μM, 5 mL) was biotinylated with Biotin-LC-NHS (100 eq.) by incubation at room temperature under gentle agitation in 50 mM HEPES, 100 mM NaCl buffer (pH=7.4). After 2 hours biotinylated hFAP was purified via PD-10 column and dialyzed overnight in HEPES buffer. The following day a StreptaWell™ (transparent 96-well) was incubated with biotinylated hFAP (100 nM, 100 μL/well) for 1 hour at room temperature and washed with PBS (3x, 200 μL/well). The protein was blocked by adding 4% Milk in PBS (200 μL/well, 30 min at RT) and then washed with PBS (3x, 200 μL/well). Immobilized hFAP was incubated for 30 minutes in the dark with serial dilutions of OncoFAP-Fluorescein (**7**) and BiOncoFAP-Fluorescein (**8**), then washed with PBS (3x, 200 μL/well). A solution of rabbit αFITC antibody (1 μg/mL, Bio-Rad 4510-7804) in 2% Milk-PBS was added to each well (100 μL/well) and incubated for additional 30 minutes in the dark. The resulting complex was washed with PBS (3x, 200 μL/well) and incubated for additional 30 minutes of protein A-HRP (1 μg/mL in 2% Milk-PBS, 100 μL/well). Each well was washed with PBS 0.1% Tween (3x, 200 μL/ well) and with PBS (3x, 200 μL/well). The substrate (TMB - 3,3’,5,5’-Tetramethylbenzidine) was added (100 μL /well) and developed in the dark for 2 minutes. The reaction was stopped by adding 50 μL of 1M sulphuric acid. The absorbance was measured at 450 nm (ref 620-650 nm) with a TECAN spark

### Internalization Studies by Confocal Microscopy Analysis

SK-RC-52.hFAP and HT-1080.hFAP cells were seeded into 4-well coverslip chamber plates (Sarstedt, Inc.) at a density of 10^4^ cells per well in RPMI or DMEM medium, respectively (1 mL, Invitrogen) supplemented with 10% Fetal Bovine Serum (FBS, Thermofisher), Antibiotic-Antimytotic (Gibco), and 10 mM HEPES (VWR). Cells were allowed to grow overnight under standard culture conditions. The culture medium was replaced with fresh medium containing the suitable Alexa488-conjugated probes (100 nM) and Hoechst 33342 nuclear dye (Invitrogen, 1 μg/mL). Colonies were randomly selected and imaged 30 min after incubation on a SP8 confocal microscope equipped with an AOBS device (Leica Microsystems).

### Animal studies

All animal experiments were conducted in accordance with Swiss animal welfare laws and regulations under the license number ZH006/2021 granted by the Veterinäramt des Kantons Zürich.

### Implantation of subcutaneous tumours

Tumour cells were grown to 80% confluence in Dulbecco’s Modified Eagle Medium (DMEM, Gibco) or RPMI-1640 (Gibco) with 10% fetal bovine serum (FBS) (Gibco) and 1% antibiotic-antimycotic (Gibco) and detached with Trypsin-EDTA 0.05%. Tumour cells were resuspended in Hanks’ Balanced Salt Solution medium. Aliquots of 5 × 10^6^ cells (100 μL of suspension) were injected subcutaneously in the flank of female athymic Balb/c AnNRj-Foxn1 mice (6 to 8 wk of age, Janvier).

### Quantitative Biodistribution of ^177^Lu-OncoFAP and ^177^Lu-BiOncoFAP in Tumour-Bearing Mice

OncoFAP-DOTAGA (**1**) and BiOncoFAP-DOTAGA (**4**) were radiolabeled with ^177^Lu (as described in the **Supplemental material**). Tumours were allowed to grow to an average volume of 500 mm^3^. Mice were randomized (n = 4/5 per group) and injected intravenously with radiolabeled preparations of ^177^Lu-OncoFAP and ^177^Lu-BiOncoFAP (250 nmol/kg; 50 MBq/kg). Mice were euthanized at different time-points (1h, 4h, 17h and 24h) after the intravenous injection by CO_2_ asphyxiation. Tumours, organs, and blood were harvested, weighted, and radioactivity was measured with a Packard Cobra Gamma Counter. Values are expressed as percent ID/g ± SD [**Figure 3**]. The %ID/g in the tumours was corrected by tumour growth rate.(*30*)

### Therapy studies with ^177^Lu-OncoFAP and ^177^Lu-BiOncoFAP in Tumour-Bearing Mice

The anti-cancer efficacy of ^177^Lu-OncoFAP and ^177^Lu-BiOncoFAP was assessed in athymic Balb/c AnNRj-Foxn1 mice bearing HT-1080.hFAP (right flank) and HT-1080.wt (wild type, left flank). ^177^Lu-OncoFAP or ^177^Lu-BiOncoFAP were intravenously administered at a dose of 250 nmol/kg, 3.7 GBq/kg (single administration; ^177^Lu-BiOncoFAP was administered on day 11, ^177^Lu-OncoFAP was administered on day 12). Therapy experiments started when the average volume of established tumours had reached 150 mm^3^. Body weight of the animals and tumour volume were daily measured and recorded. Tumour dimensions were measured with an electronic caliper and tumour volume was calculated with the formula (long side, mm) × (short side, mm) × (short side, mm) × 0.5. Animals were euthanized when one or more termination criterium indicated by the experimental license was reached (e.g., weight loss > 15%). Prism 7 software (GraphPad Software) was used for data analysis.

## Results

### Preparation of OncoFAP and BiOncoFAP conjugates

The dimeric ligand (BiOncoFAP-COOH, compound **13**) was chemically synthesized exploiting L-Lysine for the multimerization of the OncoFAP targeting moiety. The free carboxylic acid served as functional group for the conjugation of fluorophores (BiOncoFAP-Fluorescein **8**, BiOncoFAP-Alexa488 **10** and BiOncoFAP-IRDye750 **12**) and of DOTAGA chelator (compound **4**). All compounds were produced in high yields and purities [**Supplemental material**]. Monovalent OncoFAP and corresponding conjugates (OncoFAP-Fluorescein **7**, OncoFAP-Alexa488 **9** and OncoFAP-IRDye750 **11**) were synthesized following established procedures.(*18*) Chemical structures of OncoFAP and BiOncoFAP derivatives are illustrated in **Figure 1** and in **Supplemental material**. Radiolabeling of OncoFAP-DOTAGA (**1**) and BiOncoFAP-DOTAGA (**4**) with ^177^Lu was achieved in high yield and purity [**Supplemental material**]. After radiolabeling, ^177^Lu-OncoFAP and ^177^Lu-BiOncoFAP retained the ability to form stable complexes with recombinant human FAP, as assessed by PD-10 co-elution experiment. Both compounds are highly hydrophilic, with experimental Log*D*_7.4_ values of – 4.02 ± 0.22 (n = 5) and – 3.60 ± 0.31 (n = 5), respectively [**Supplemental material**].

**Figure 1.**
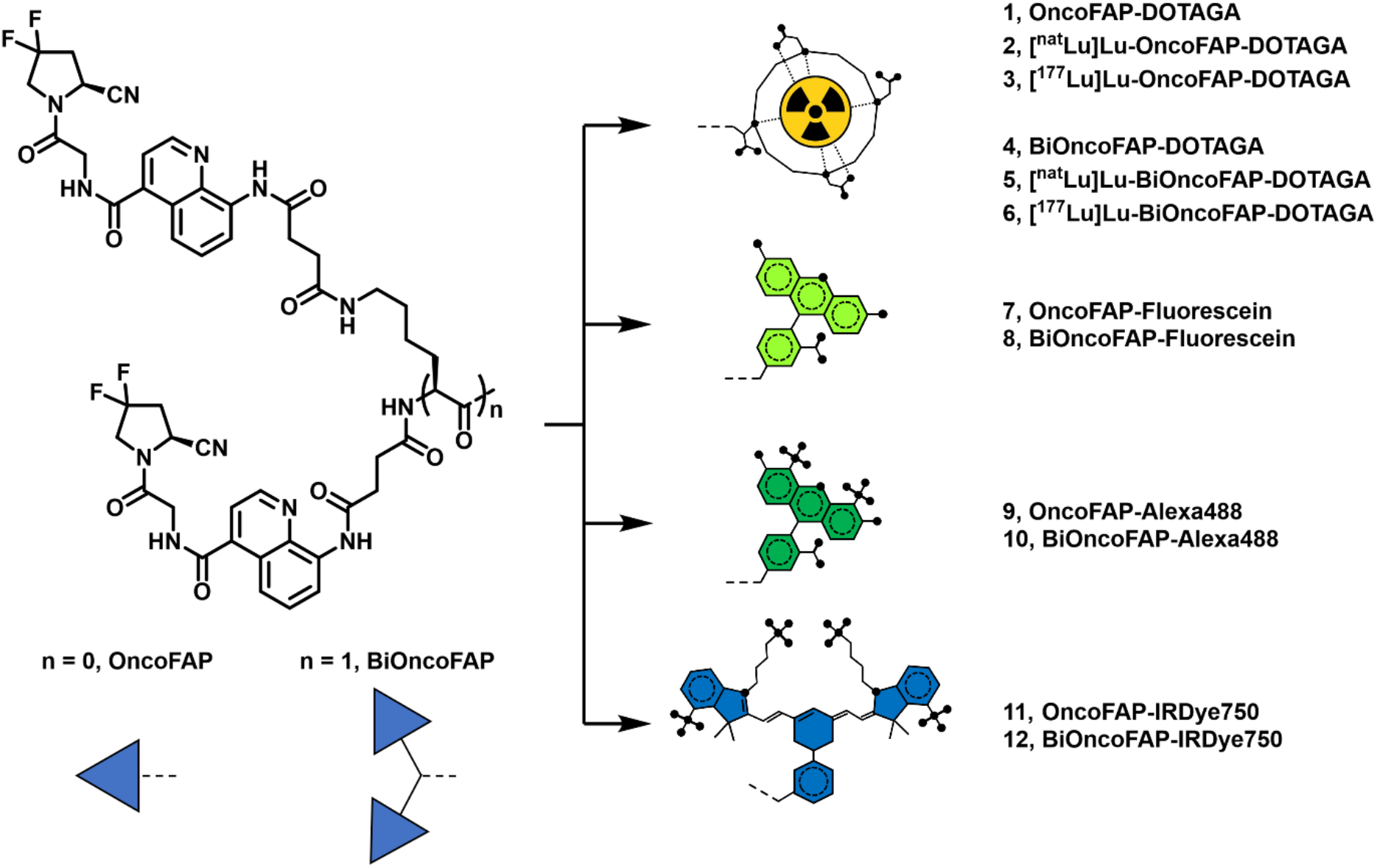
Chemical structures of OncoFAP and BiOncoFAP conjugates (schemes). BiOncoFAP and OncoFAP and their DOTAGA, Fluorescein, Alexa488 and IRDye750 conjugates are presented.

### In vitro inhibition assay against hFAP

We evaluated the inhibitory activity of OncoFAP-DOTAGA (**1**), BiOncoFAP-DOTAGA (**4**) and their ^nat^Lu cold-labeled derivatives (**2** and **5**, respectively) against human recombinant FAP (hFAP). Compounds **4** and **5** displayed enhanced inhibitory activity against the target (IC_50_ = 168 and 192 pM, respectively) compared to their monovalent counterparts (OncoFAP-DOTAGA IC_50_ = 399 pM, [^nat^Lu]Lu-OncoFAP-DOTAGA IC_50_ = 456 pM) [**Figure 2A and 2B**].

**Figure 2.**
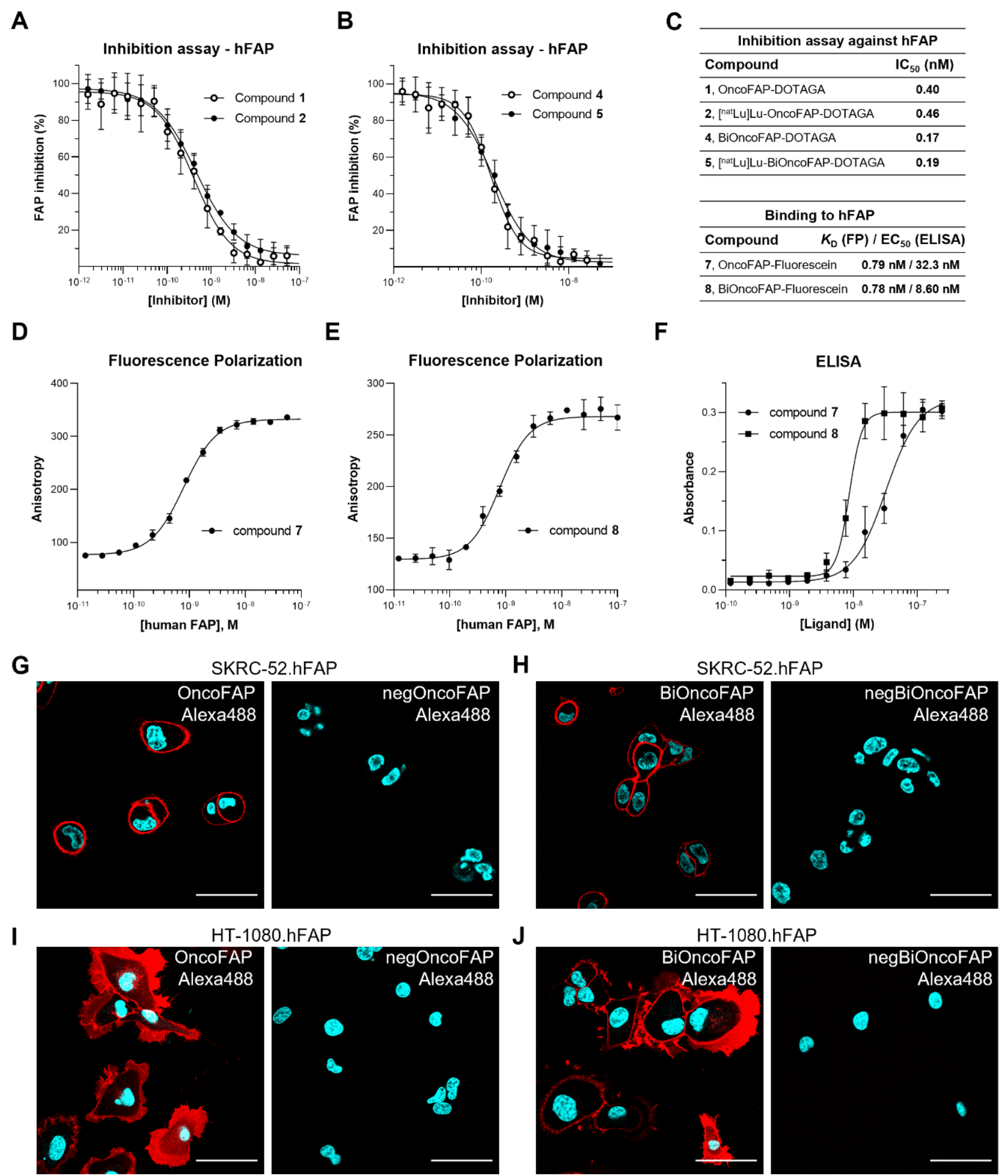
Enzymatic assays performed with (*A*) OncoFAP-DOTAGA, (*B*) BiOncoFAP-DOTAGA, and their corresponding cold ^nat^Lu-labeled derivatives (**2** and **5**). *(C)* Binding affinity and IC50 values of OncoFAP and BiOncoFAP derivatives towards hFAP. Affinity measurement of (*D*) OncoFAP-Fluorescein (**7**) and (*E*) BiOncoFAP-Fluorescein (**8**) to recombinant human Fibroblast Activation Protein by fluorescence polarization. Both compounds showed ultra-high affinity for the FAP target. (*F*) ELISA experiment on OncoFAP-Fluorescein (**7**) and BiOncoFAP-Fluorescein (**8**) against hFAP. Confocal microscopy images after incubation of OncoFAP-Alexa488 (**9**) and BiOncoFAP-Alexa488 (**10**) with SK-RC-52.hFAP (*G* and *H*, respectively) or HT-1080.hFAP to (*I* and *J*, respectively). Red = fluorescein derivatives staining; blue = Hoechst 33342 staining. (Scale bar = 50 μm.).

**Figure 3.**
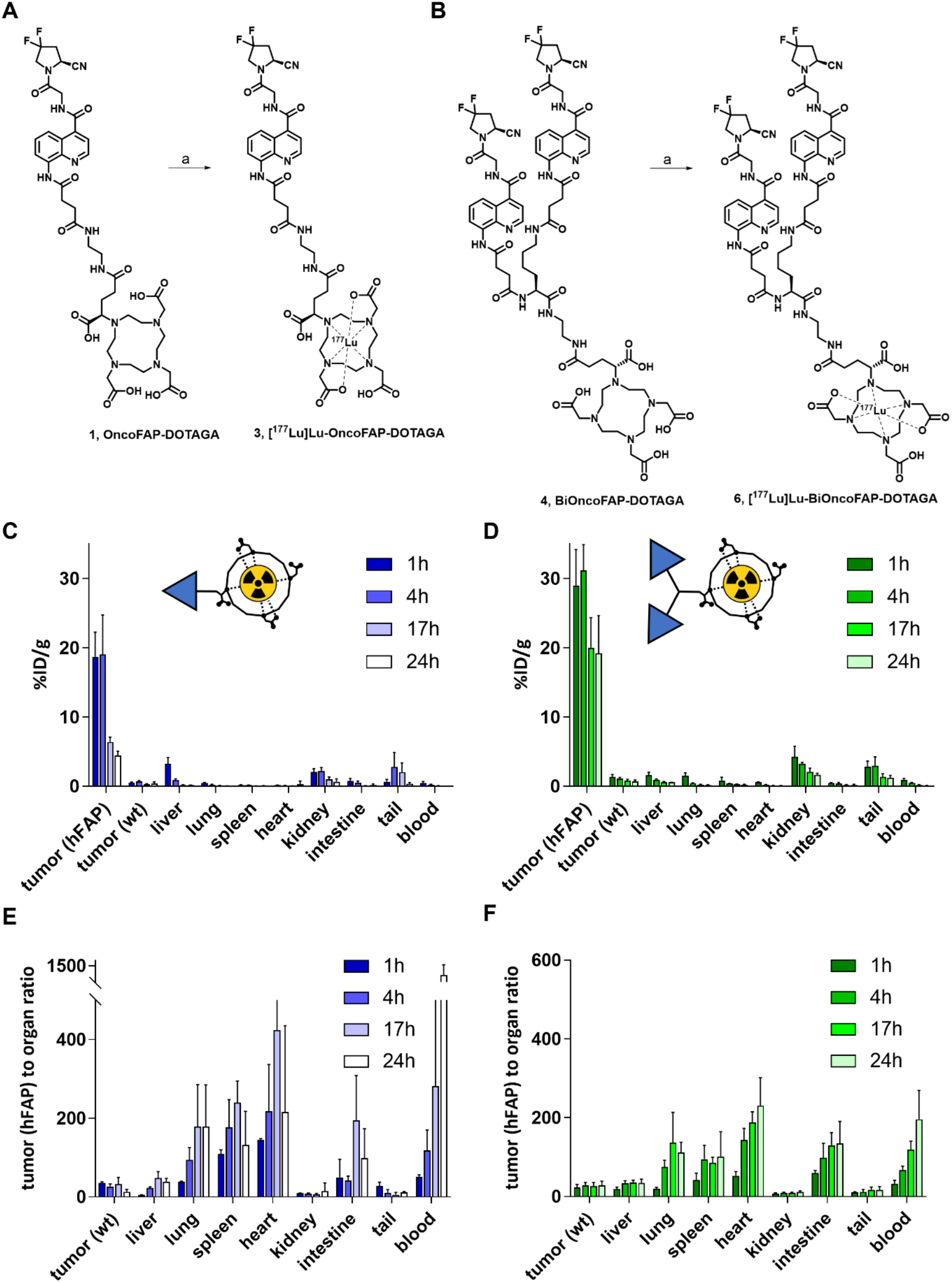
Radiolabeling scheme of (*A*) OncoFAP-DOTAGA (**1**) and (*B*) BiOncoFAP-DOTAGA (**4**) with lutetium-177 to obtain ^177^Lu-OncoFAP (**3**) and ^177^Lu-BiOncoFAP (**6**). Reagents and conditions: a) ^177^Lu, acetate buffer, 90°C, 15 min. Quantitative *in vivo* biodistribution and tumour-to-organ ratio of (*C, E*) ^177^Lu-OncoFAP and (*D, F*) ^177^Lu-BiOncoFAP at different time points after intravenous administration (250 nmol/kg, 50 MBq/kg) in mice bearing HT-1080.wt and HT-1080.hFAP tumours.

### Assessment of binding properties of BiOncoFAP to soluble and immobilized human. recombinant Fibroblast Activation Protein

In order to study the binding properties of OncoFAP and BiOncoFAP to soluble hFAP, we measured the affinity constant (*K*_D_) of the corresponding fluorescein conjugates (compound **8**, OncoFAP-Fluorescein and compound **9**, BiOncoFAP-Fluorescein) in Fluorescence Polarization assays [**Figure 2D** and **2E**]. Compounds **8** and **9** exhibited comparable sub-nanomolar *K*_D_ values against hFAP (respectively 795 and 781 pM). Moreover, both compounds showed to be very selective for FAP and did not bind to a set of non-target proteins up to micromolar concentrations [**Figure S2, Supplemental material**]. Our data confirms that the dimerization does not impair the affinity and selectivity of BiOncoFAP for its target. Then, we studied the binding affinity to hFAP immobilized on a solid support of the dimeric ligand. In a comparative ELISA, BiOncoFAP-Fluorescein exhibited a lower *K*_D_ compared to OncoFAP-Fluorescein (8.60 nM vs 32.3 nM, respectively) [**Figure 2C, 2F**].

### Confocal microscopy analysis on tumour cells

Binding of BiOncoFAP to FAP-positive SK-RC-52 and HT-1080 cancer cells and internalization were assessed by confocal microscopy analysis using the corresponding Alexa-488 conjugate (compound **10**). OncoFAP-Alexa488 (compound **9**) was used in the same experiment positive control, while untargeted analogues were included as non-binding negative controls [chemical structures are depicted in the **Supplemental material**]. OncoFAP and BiOncoFAP displayed comparable binding features on living tumour cells. Both compounds showed lack of internalization on FAP-positive SK-RC-52 cells, while high membrane trafficking was observed when compounds were incubated on HT-1080.hFAP cells [**Figure 2G** and **2J**].

### Stability studies

The stability of cold-labeled [^nat^Lu]Lu-BiOncoFAP-DOTAGA was assessed in human and mouse serum after incubation at 37°C for 24, 48, 72 and 120 hours. The test compound exhibited half-life longer than 5 days in all experimental conditions. No loss of lutetium (^nat^Lu) from the DOTAGA chelator was detected [**Supplemental material, Figure S3**].

### Biodistribution of OncoFAP and BiOncoFAP in tumour-bearing mice

Qualitative biodistribution of OncoFAP and BiOncoFAP was assessed in tumour-bearing mice using a near-infrared fluorophore (IRDye750) as detection agent. Macroscopic imaging of mice implanted with SK-RC-52.hFAP (right flank) and SK-RC-52.wt (left flank) tumours revealed that both OncoFAP-IRDye750 (compound **11**) and BiOncoFAP-IRDye750 (compound **12**) selectively accumulated in FAP-positive tumours [**Figure S4-S5, Supplemental material**]. Interestingly, BiOncoFAP-IRDye750 conjugate exhibited a longer residence time at the site of disease. Encouraged by these results, we studied the quantitative biodistribution of ^177^Lu-BiOncoFAP in athymic Balb/c mice bearing HT-1080.hFAP (right flank) and HT-1080.wt (left flank) tumours. A direct comparison with ^177^Lu-OncoFAP was included in the experiment [**Figure 3**]. Both compounds accumulated selectively in FAP-positive tumours shortly after intravenous administration. The dimeric ^177^Lu-BiOncoFAP product exhibited a more stable and prolonged tumour uptake as compared to its monovalent counterpart (∼20% ID/g vs ∼4% ID/g, 24h after systemic administration). Notably, ^177^Lu-BiOncoFAP did not show significant uptake in healthy organs, with favorable tumour-to-organ ratio (e.g., 12-to-1 tumour-to-kidney and 34-to-1 tumour-to-liver ratio, at the 24 h time point).

### Therapy study

Therapeutic efficacy of ^177^Lu-OncoFAP and of ^177^Lu-BiOncoFAP was assessed in mice bearing HT-1080.hFAP tumours on the right flank and HT-1080.wt tumours on the left flank [**Figure 4**]. Systemic administration of both compounds at therapeutic doses (70 MBq/mouse, 250 nmol/Kg) resulted in selective and potent anti-cancer activity against the growth of HT-1080.hFAP as compared to mice injected with saline. The most active compound in our therapy study was ^177^Lu-BiOncoFAP. Tumour growth of FAP-negative lesions (HT-1080.wt) was not influenced by the treatment with ^177^Lu-OncoFAP and with ^177^Lu-BiOncoFAP.

**Figure 4.**
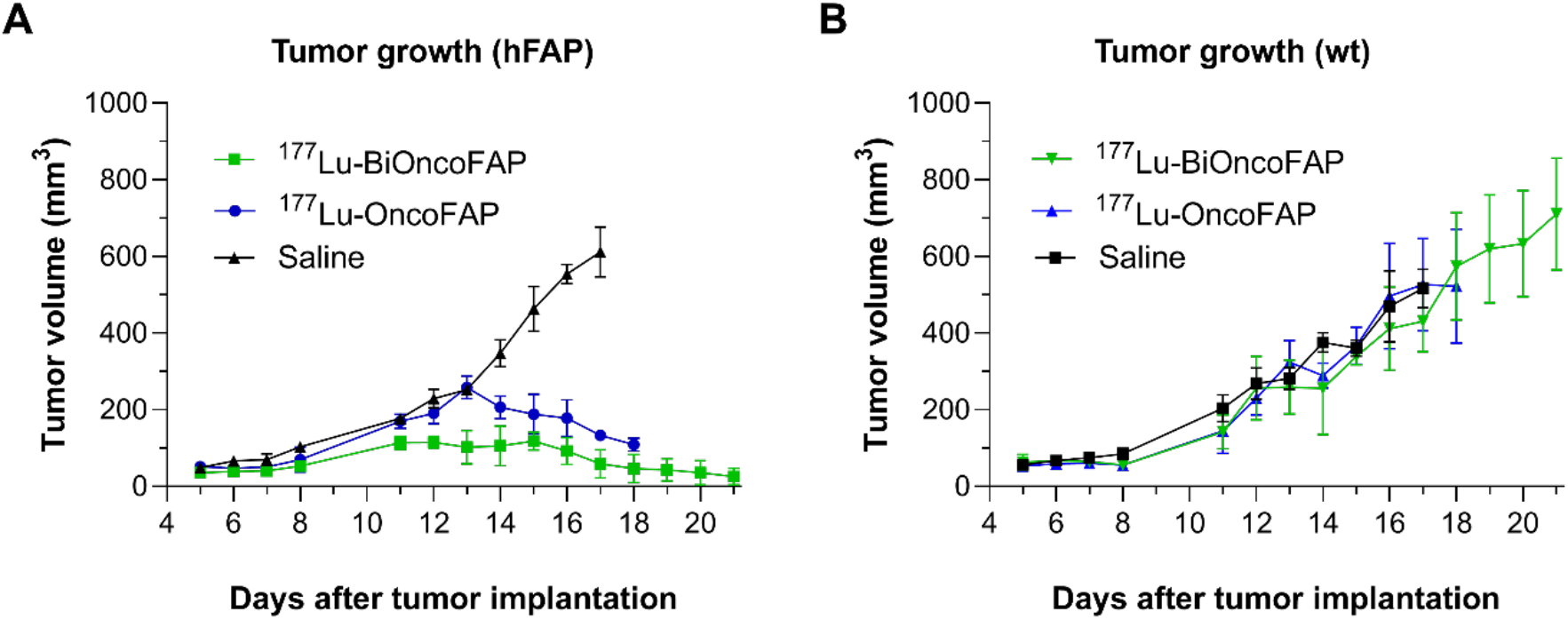
Therapeutic activity after a single administration (250 nmol/kg, 70 MBq/mouse) of ^177^Lu-OncoFAP (compound **3**) and ^177^Lu-BiOncoFAP (compound **6**) in Balb/c nu/nu mice bearing HT-1080.hFAP tumour in the right flank (*A*) and HT-1080.wt tumour in the left flank (*B*). The efficacy of the different treatments was assessed by daily measurement of tumour volume (mm^3^) after administration of the different compounds. Data points represent mean tumour volume ± SEM.

## Discussion

FAP-targeted radiopharmaceuticals may revolutionize the field of radioligand imaging of cancer, because of their applicability to many types of malignancies and for the excellent tumour selectivity which has already been proven at the clinical level.(*12,13,19*) Other SMRC products, ^177^Lu-PSMA-617 and Lutathera®, are limited to certain specific cancer indications and may be taken up by certain normal organ structures.(*31,32*) FAP is mainly expressed in the stroma of solid malignancies, thus adding a new element of differentiation compared to previously established targeting platforms, based on SSTR-2 and PSMA as cellular antigens.(*12,13,16,17*) In this context, accurate selection of the radionuclide payload is crucial for the success of FAP-targeted radiotherapy. While alpha-emitters are characterized by a short range, typically >100 μm (*33*) which may be insufficient, the use of a beta-emitter radionuclide such as ^177^Lu (path length of ∼1.5 mm) (*33*) should enable the killing not only of the stromal cells, but also of surrounding tumour cells.(*34,35*)

Sustained accumulation of SMRCs in tumours is fundamental for the effective delivery of high radiation doses over time at the site of disease, and therefore for the success of the therapeutic treatment. Among different approaches employed in the past, the dimerization of high affinity ligands has been proposed as a strategy to enhance residence time in antigen-positive structures (i.e., in FAP-positive tumours).(*23,36*–*39*) Dimeric ligands present higher chances of re-binding to their target, with slower off-rates as compared to their monovalent counterparts (*40*) However, increase in the binding valency typically leads to higher uptake in healthy tissues.(*21,38,39*) To the best of our knowledge, two dimeric FAP-targeting radionuclides named DOTA/DOTAGA.(SA.FAPi)_2_ (*41,42*) and DOTA-2P(FAPi)_2_ (*21*) have recently been described. While preclinical biodistribution data are not available for DOTA/DOTAGA.(SA.FAPi)_2_, DOTA-2P(FAPi)_2_ was extensively characterized in HCC-PDX-1 mouse model. Despite its slightly increased tumour uptake compared to FAPi-46 (∼9% vs ∼4% ID/g 1h p.i.), the dimeric ligand presents low tumour-to-organ ratios both at 1 and 4h, with a particular liability for the kidney (∼1.2-to-1 and ∼1.5-to-1 tumour-to-kidney ratio, respectively).(*21*) BiOncoFAP, the novel homodimeric FAP-targeting small organic ligand described in this article, shows specific and persistent tumour uptake (∼30% ID/g 1 h p.i. and ∼20% ID/g 24 h p.i.) in HT-1080.hFAP tumour bearing mice. Remarkably, ^177^Lu-BiOncoFAP presents a clean preclinical biodistribution profile with high tumour-to-organ ratios even at early time-points (e.g., ∼7-to-1 and ∼10-to-1 tumour-to-kidney, and ∼20-to-1 and ∼34-to-1 tumour-to-liver ratio, at the 1h and 4h time points, respectively).

The *in vivo* anti-cancer activity of [^177^Lu]Lu-FAPI-46 (beta-emitter) and [^225^Ac]Ac-FAPI-46 (alpha-emitter) has been recently evaluated in PANC-1 tumour bearing mice, a xenograft model of pancreatic cancer characterized by high stromal expression of FAP.(*35*) Both products showed only a limited tumour growth suppression even at the highest dose (i.e., 30 kBq/mouse for [^225^Ac]Ac-FAPI-46 and 30 MBq/mouse for [^177^Lu]Lu-FAPI-46).

Collectively, our biodistribution and therapy results show that both ^177^Lu-OncoFAP and ^177^Lu-BiOncoFAP are able to efficiently localize at the tumour site and produce potent anti-cancer effect in mice bearing subcutaneous FAP-positive tumours, after a single administration at a dose of 70 MBq/mouse (∼2 mCi/mouse). Compared to the monomeric ^177^Lu-OncoFAP, our new bivalent ^177^Lu-BiOncoFAP displayed an enhanced *in vivo* anti-tumour activity. As expected, lack of tumour suppression was observed for the FAP-negative tumours (HT-1080.wt), which were used as internal control to appreciate the specificity of OncoFAP-based theranostic products towards FAP-positive solid lesions.

Considering exquisite selectivity for cancer lesions and pan-tumoural properties of FAP-target radioligand therapeutics, this new class of radiopharmaceutical products may represent a breakthrough in cancer therapy.(*12*) Interim reports on the efficacy of FAP-targeting peptides and small organic ligands developed so far have shown limitation of this therapeutic strategy.(*26,43*) Escalation of the dose of radiolabeled FAP-targeting peptides is limited by their intrinsically high kidney uptake at late time-points.(*43*–*45*) Therapy with small organic ligands based on FAPI-46 may be limited by their short residence time in the tumour.(*26,46*) We have developed ^177^Lu-BiOncoFAP, a new radioligand therapeutic product with prolonged *in vivo* tumour uptake, and highly favorable tumour-to-kidney ratios. Future clinical studies in a basket of indications will provide clarity on the therapeutic efficacy of this novel FAP-targeted product.

## Conclusion

_177_Lu-BiOncoFAP is a promising FAP-targeted SMRC product for tumour therapy. This novel bivalent FAP-targeted compound binds its target with high affinity and shows long residence time in tumour lesions, with favorable tumour-to-organ ratios. Once administered at therapeutic doses, ^177^Lu-BiOncoFAP potently inhibits growth of FAP-positive tumours in mice. Our data support clinical development of ^177^Lu-BiOncoFAP in the frame of targeted radioligand therapy.

## Acknowledgements

The authors would like to thank Ettore Gilardoni for performing exact mass analysis of compounds presented in this article, and Frederik Peissert and Luca Prati for their support with small molecule-ELISA experiments.

## Availability of data and material

Additional data is available in the supplement material

