## Supplemental material for "A novel dimeric FAP-targeting small molecule-radio conjugate with high and prolonged tumour uptake"

### Index

Quality control of radiosynthesis - Coelution Experiments of Ligand–Protein Complexes. 26

### Chemistry

#### Material and methods

Liquid Chromatography-Mass Spectrometry (LC-MS) spectra presented were recorded on an Agilent 6100 Series Single Quadrupole MS system combined with an Agilent 1200 Series LC, using an InfinityLab Poroshell 120 EC-C18 Column, 2.7 µm, 4.6 × 50 mm at a flow rate of 0.8 mL/min, 10% ACN in 0.1% aq. HCOOH to 100% ACN in 3 or 10 min.

Reversed-phase high-pressure liquid chromatography (RP-HPLC) were performed on an Agilent 1200 Series RP-HPLC with PDA UV detector, using a Synergi 10µm, MAX-RP 80Å 10 × 250 mm C18 column at a flow rate of 5 mL/min with linear gradients of solvents A and B (A = Millipore water with 0.1% TFA, B = ACN with 0.1% TFA).

High-Resolution mass spectrometry (HR-MS) were performed on a Q Exactive Mass Spectrometer (Thermo Fisher Scientific). The analyte was injected directly into the MS at a flow rate of 4 µL/min. Both MS1 and MS2 spectra were recorded. MS1 spectra were obtained with a resolution of 70000. MS2 spectra were obtained by inducing fragmentation of the molecule with a NCE (normalized collision energy) =25 and with a resolution of 70000.

#### Chemistry and radiochemistry

(*S*)-4-((4-((2-(2-cyano-4,4-difluoropyrrolidin-1-yl)-2-oxoethyl)carbamoyl)quinolin-8-yl)amino)-4-oxobutanoic acid (named OncoFAP-COOH), OncoFAP-Fluorescein (compound **7**), OncoFAP-Alexa488 (compound **9**) and OncoFAP-IRDye750 (compound **11**) were synthesized as previously reported by Millul and co-workers.(*1*)

OncoFAP-DOTAGA (**1**) and BiOncoFAP-DOTAGA (**4**) were labeled with cold lutetium by incubation with $[^{nat}Lu]LuCl_3$ in acetate buffer at 90°C for 15 minutes to obtain $[^{nat}Lu]$Lu-OncoFAP-DOTAGA (**2**) and $[^{nat}Lu]$Lu-BiOncoFAP-DOTAGA (**5**) which were used as reference compounds for *in vitro* characterization (inhibition assay and serum stability).

Radiolabeling of OncoFAP-DOTAGA (**1**) and BiOncoFAP-DOTAGA (**4**) with lutetium-177 was performed with two different specific activities for the two different studies (biodistribution and therapy).

Prior to the biodistribution study, precursors (compound **1** or **4**, 100 nmol) were dissolved in 100 µL of PBS and diluted with 200 µL of sodium acetate (1 M in water, pH = 8). 20 MBq of $^{177}$Lu solution were added and the mixture was heated at 90°C for 15 minutes, followed by dilution with 1600 µL of PBS to achieve final volume of 2 mL.

Prior to the therapy study, precursors (compound **1** or **4**, 5 nmol) were dissolved in 5 µL of PBS, then sodium acetate buffer (30 µL, 1 M in water) and 70 MBq of $^{177}$Lu solution were added. The mixture was heated at 90°C for 15 minutes followed by dilution with 130 µL of PBS to afford a final volume of 200 µL.

Quality control of radiosynthesis was performed using radio-HPLC. The possibility to form a stable complex between the so obtained $^{177}$Lu-radiolabeled derivatives and the target antigen was tested by co-incubating the compounds with recombinant human FAP and loading the mixture onto a desalting PD-10 column run by gravity (details are reported below).

#### Synthetic schemes

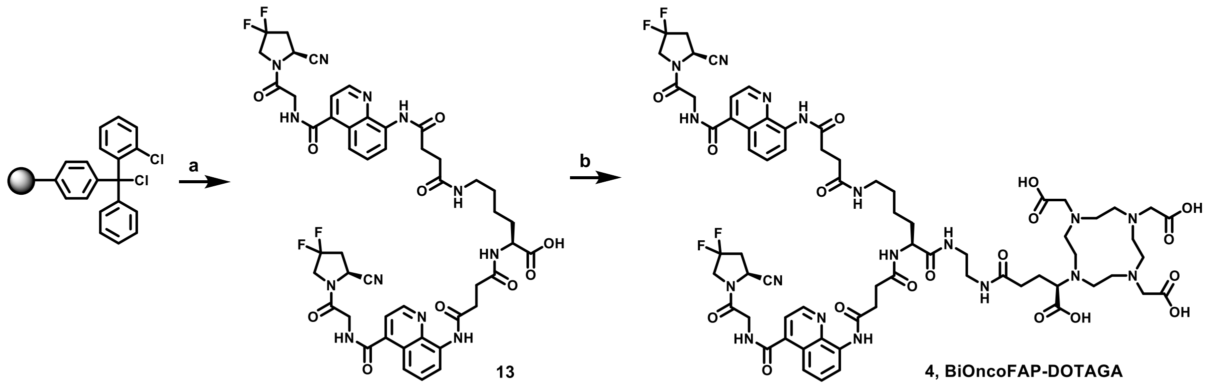

**Scheme S1**. Synthesis of BiOncoFAP-DOTAGA (**4**). Reagents and conditions: a) i) NHFmoc-L-Lys(Fmoc)-OH, NMM, dry DCM, 6h, r.t.; ii) MeOH, NMM, dry DCM, 30 min, r.t.; iii) Piperidine/DMF 20% v/v, 20 min, r.t.; iv) OncoFAP-COOH, HATU, DIPEA; DMF, 1h, r.t.; v) TFA/DCM 30% v/v, 1h, r.t.; b) i) *N*-hydroxysuccinimide, HATU, DIPEA, DMF, 30 min, r.t.; ii) (R)-DOTA-GA-$NH_2$, overnight, r.t.

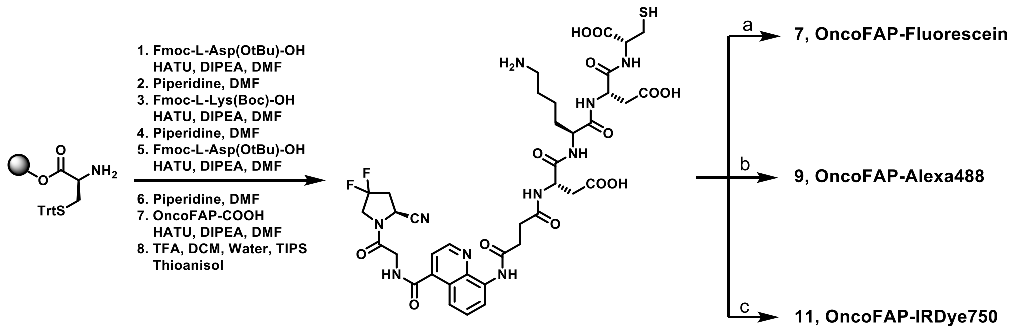

14, OncoFAP-Asp-Lys-Asp-Cys

**Scheme S2.** Synthesis of OncoFAP-Fluorescein (**7**), OncoFAP-Alexa488 (**9**) and OncoFAP-IRDye750 (**11**). Reagents and conditions: a) Maleimido-Fluorescein, DMF, PBS, 3 h, r.t.; b) AlexaFluor488-C5-Maleimide, DMSO, PBS, 3 h, r.t.; c) IRDye750-Maleimide, DMF, PBS, 3h, r.t.

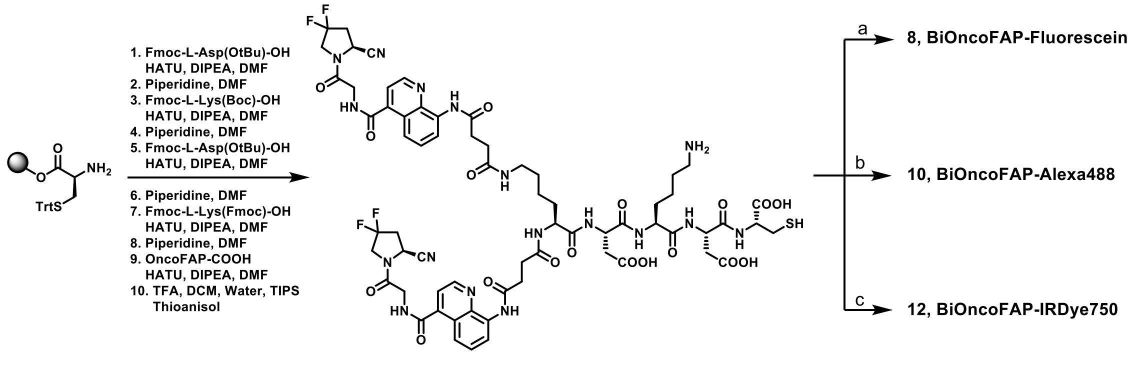

15, BiOncoFAP-Asp-Lys-Asp-Cys

**Scheme S3.** Synthesis of BiOncoFAP-Fluorescein (**8**), BiOncoFAP-Alexa488 (**10**) and BiOncoFAP-IRDye750 (**12**). Reagents and conditions: a) Maleimido-Fluo, DMF, PBS, 3 h, r.t.; b) AlexaFluor488-C5-Maleimide, DMSO, PBS, 3 h, r.t.; c) IRDye750-Maleimide, DMF, PBS, 3h, r.t.

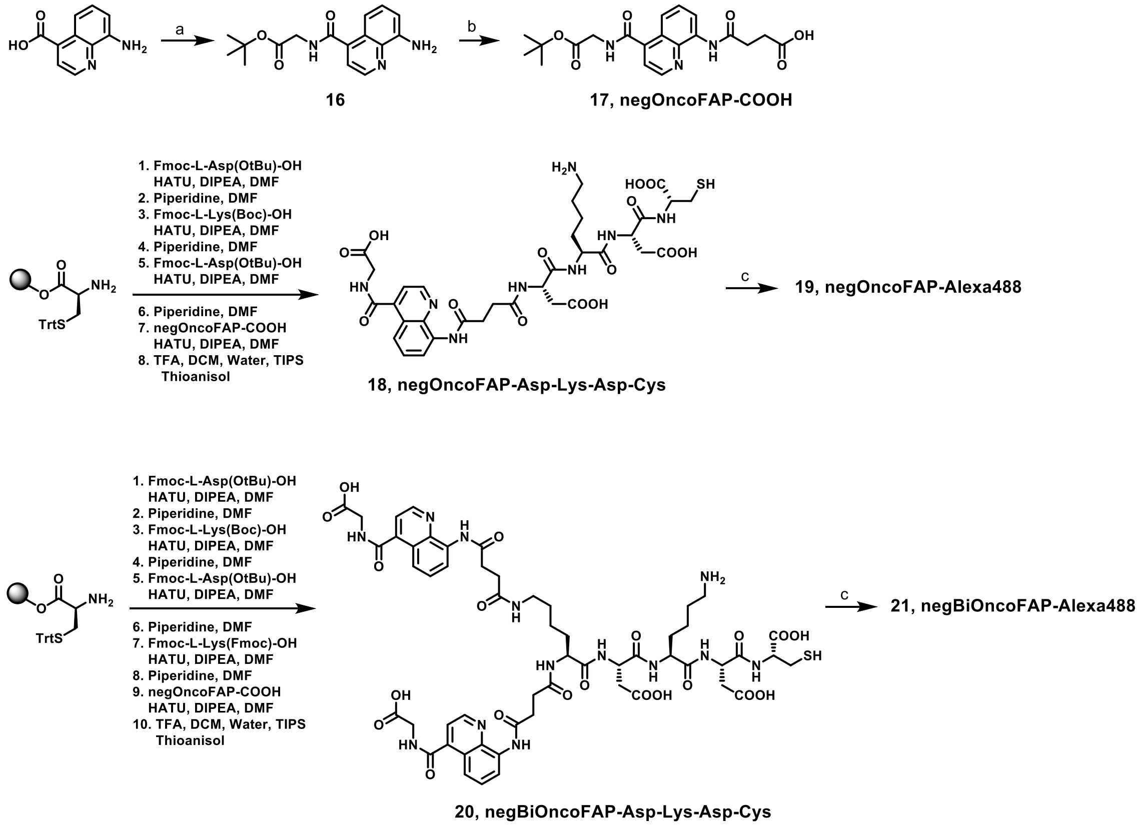

**Scheme S4.** Synthesis of negOncoFAP-Alexa488 (**19**) and negBiOncoFAP-Alexa488 (**21**). Reagents and conditions: a) Gly-OtBu*HCl, HATU, DIPEA, DCM/DMF, 30 min, 0°C to r.t.; b) Succinic anhydride, DMAP, THF, 1 h, 55°C; c) AlexaFluor488-C5-Maleimide, DMSO, PBS, 3 h, r.t.

**Synthesis of BiOncoFAP-COOH (13)**

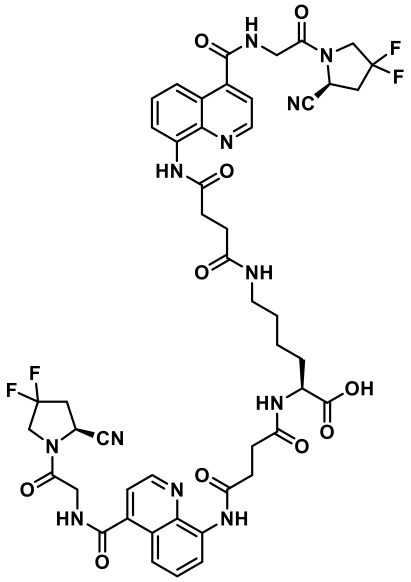

To a solid-phase synthesis syringe, 2-chlorotrityl resin (300 mg) was added and then swollen with dry DCM for 15 min. Fmoc-L-Lys(Fmoc)-OH (89 mg, 0.15 mmol, 1 eq.) and 4-Methylmorpholine (45 µL, 0.40 mmol, 2.7 eq.) were sequentially added to the resin and the mixture was allowed to react for 3 h. Next, a capping step with methanol / 4-Methylmorpholine / DCM (1:2:7 ratio, 5 mL, 30 min) was carried out, following by a wash with DMF and Fmoc-removal with 20% solution of Piperidine in DMF (10 mL). The resin was then treated with a solution of OncoFAP-COOH (137 mg, 0.300 mmol, 2.0 eq.), HATU (86 mg, 0.22 mmol, 1.5 eq.) and DIPEA (97 µL, 0.75 mmol, 5.0 eq.) in DMF (5 mL) for 1 h. After multiple washing with DMF, the resin was submitted to the cleavage with 30% solution of TFA in DCM (10 mL) for 1 h. The cleaved solution was recovered, concentrated under *vacuo* and purified by Reverse Phase Flash-Chromatography (gradient: water/acetonitrile + 0.1% FA 98:2 to 0:100 in 45 min). The fractions were collected and lyophilized to afford a white solid (30 mg, 0.029 mmol, 19% yield). MS(ES+) m/z 1029.3 $(M+H)^+$

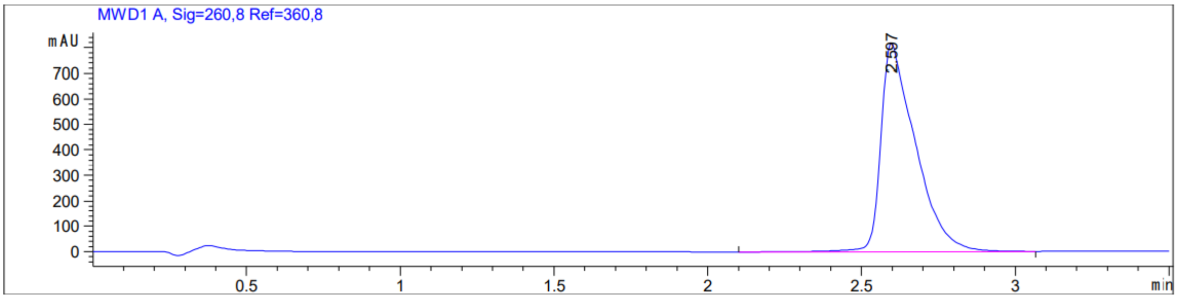

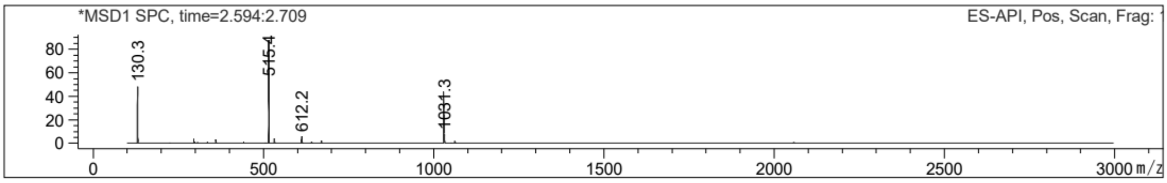

##### Synthesis of BiOncoFAP-DOTAGA (4)

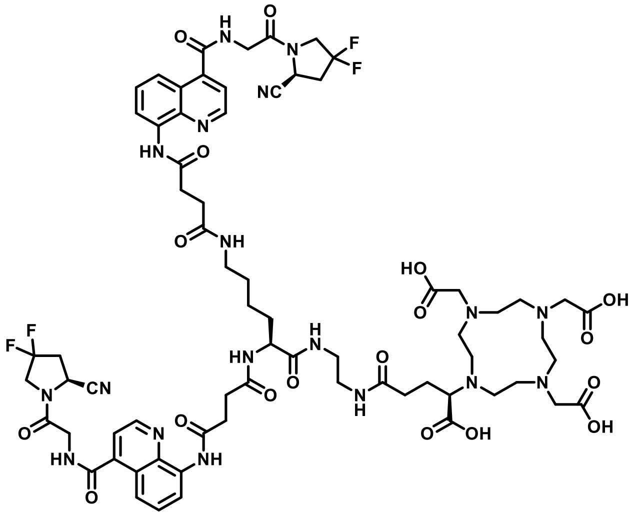

*N*-Hydroxysuccinimide (40 mg, 0.35 mmol, 2 eq.), HATU (80 mg, 0.21 mmol, 1.2 eq.) and DIPEA (0.15 mL, 0.88 mmol, 5 eq.) were added to a solution of BiOncoFAP-COOH (180 mg, 0.175 mmol, 1 eq.) in dry DMF (5 mL). The reaction solution was stirred for 30 minutes at room temperature, then (*R*)-DOTA-GA-$NH_2$ (180 mg, 0.35 mmol, 2 eq.) was added. The resulting mixture was stirred vigorously overnight, then diluted with milliQ water (5 mL) and purified via RP-HPLC (90:10 to 0:100 ACN/water + 0.1% TFA in 12 min). The desired fractions were collected and lyophilized to afford a white solid. (140 mg, 52%).

<u>HPLC-UV:</u>

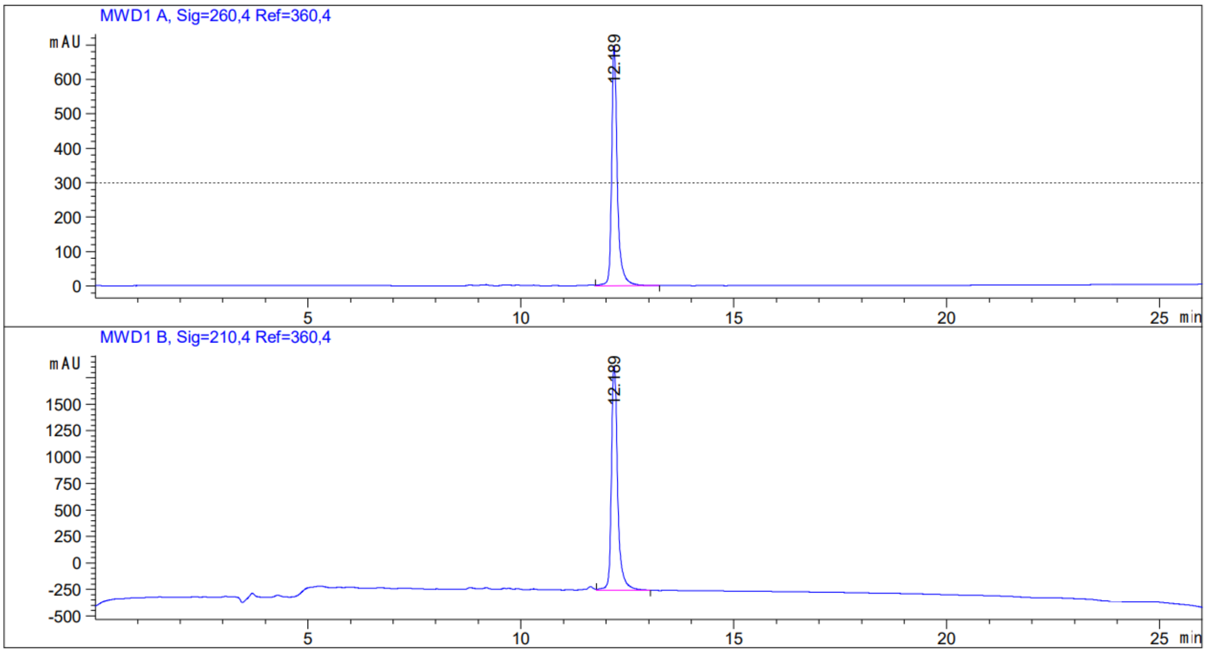

LC-UV/MS:

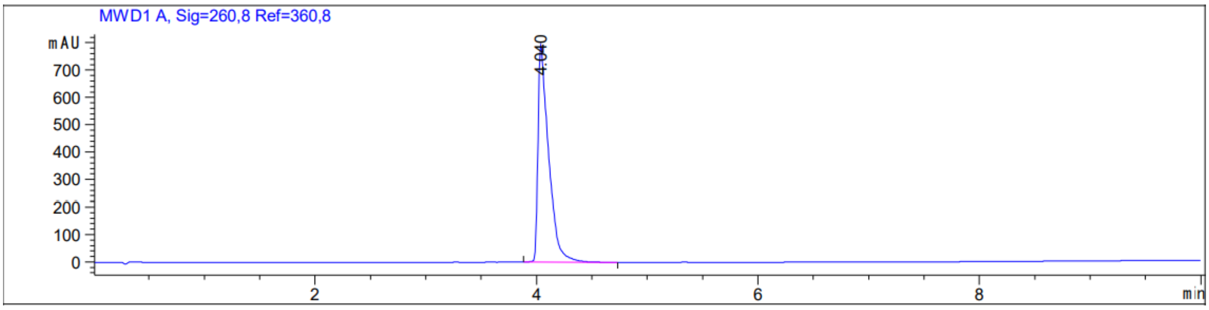

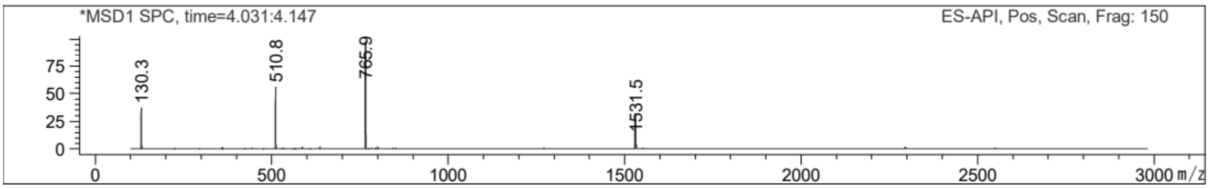

HR-MS. Predicted M/z = 1529.62198; Experimental M/z = 1529.6222; Accuracy = 0.14 ppm

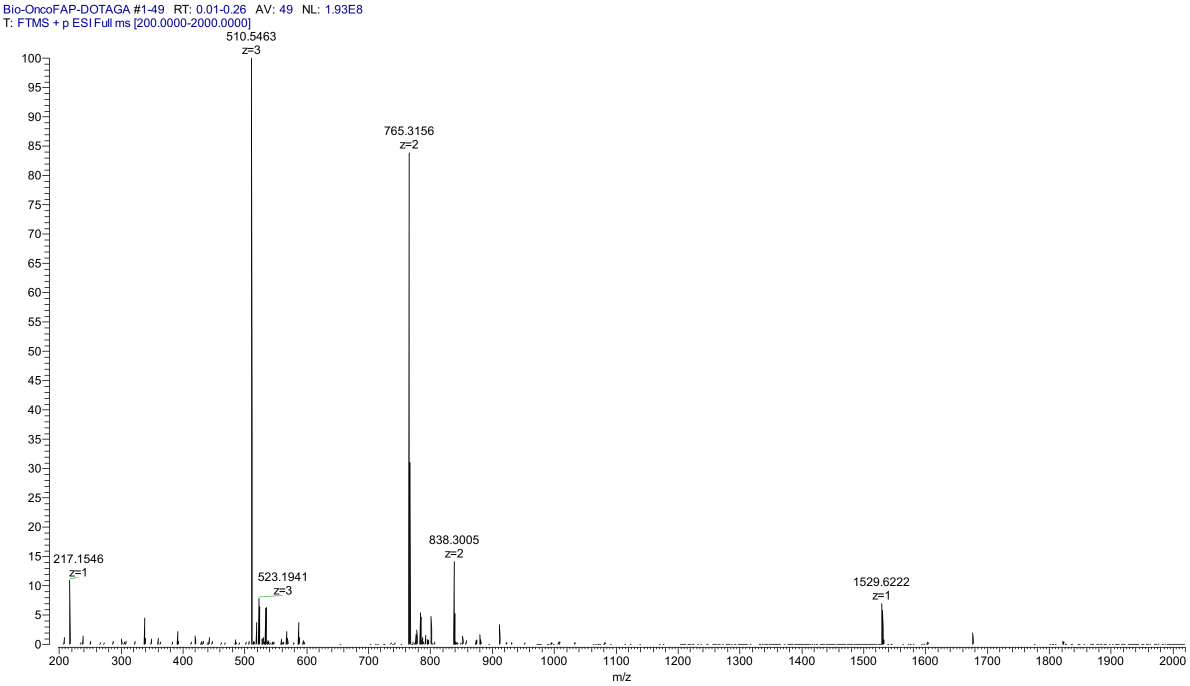

#### Synthesis of [$^{nat}$Lu]Lu-OncoFAP-DOTAGA (2)

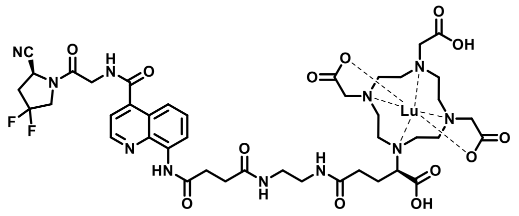

To a solution of OncoFAP-DOTAGA (compound **1**, 0.96 mg, 1 µmol, 1 eq.) in 300 µL acetate buffer (aqueous solution, 1 M, pH 8), a freshly prepared solution of $LuCl_3$ hexahydrate (0.78 mg, 2 µmol, 2 eq.) in 0.05N HCl (1.5 mL) was added. The resulting mixture was stirred at 95°C for 10-15 minutes, then purified via RP-HPLC (90:10 to 0:100 ACN/water + 0.1% TFA in 12 min). The desired fractions were collected and lyophilized to afford a white solid. (0.8 mg, 71%)

<u>LC-MS</u>:

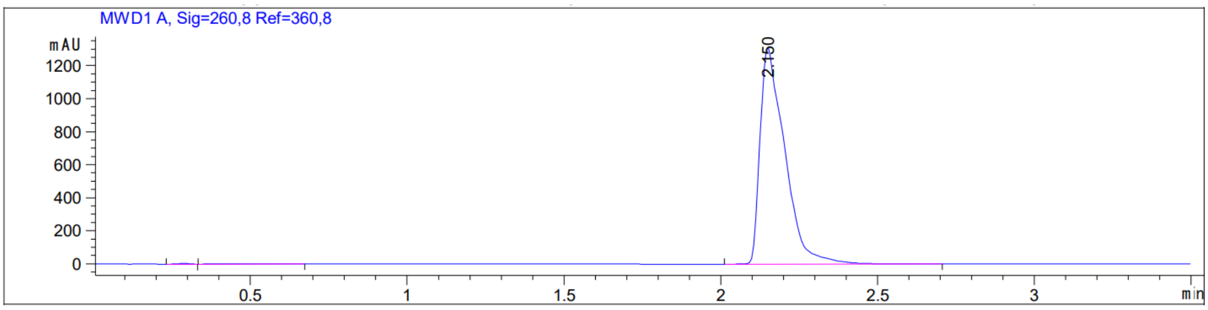

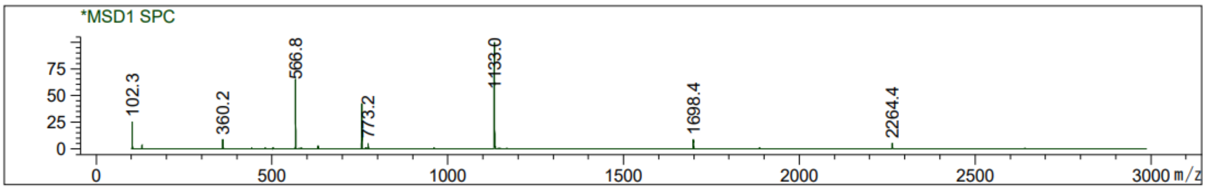

HR-MS: Predicted M/z = 1132.31946; Experimental M/z = 1132.32056; Accuracy = 0.97 ppm

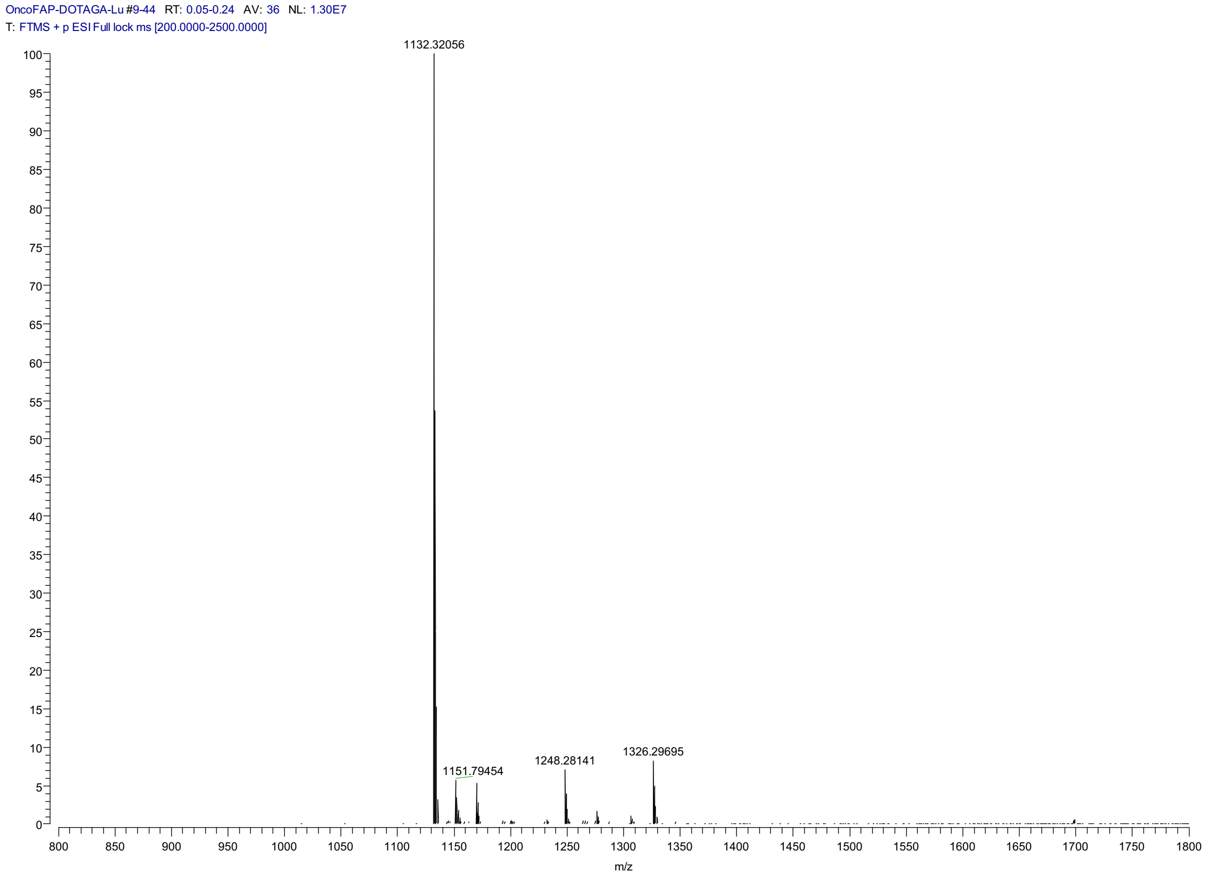

##### Synthesis of [$^{nat}$Lu]Lu-BiOncoFAP-DOTAGA (5)

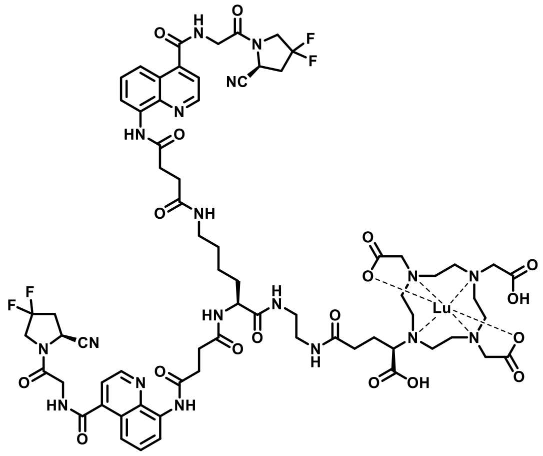

To a solution of BiOncoFAP-DOTAGA (compound **4**, 1.5 mg, 1 µmol, 1 eq.) in 300 µL acetate buffer (aqueous solution, 1 M, pH 8), a freshly prepared solution of $LuCl_3$ hexahydrate (0.78 mg, 2 µmol, 2 eq.) in 0.05N HCl (1.5 mL) was added. The resulting mixture was stirred at 95°C for 10-15 minutes, then purified via RP-HPLC (90:10 to 0:100 ACN/water + 0.1% TFA in 12 min). The desired fractions were collected and lyophilized to afford a white solid. (1.2 mg, 67%)

LC-MS:

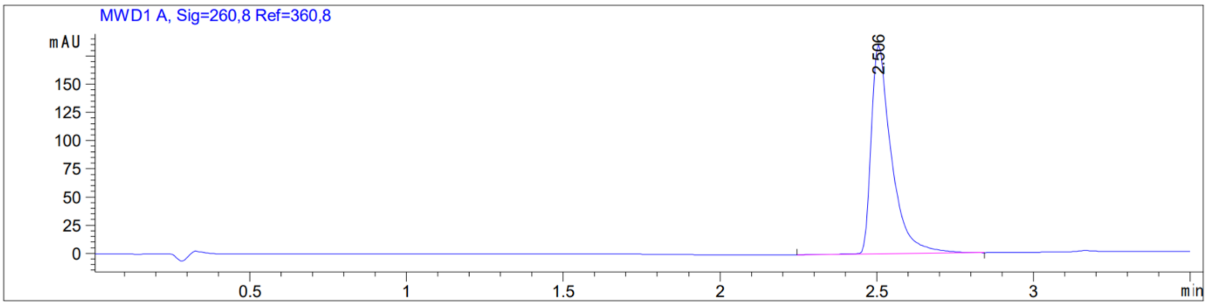

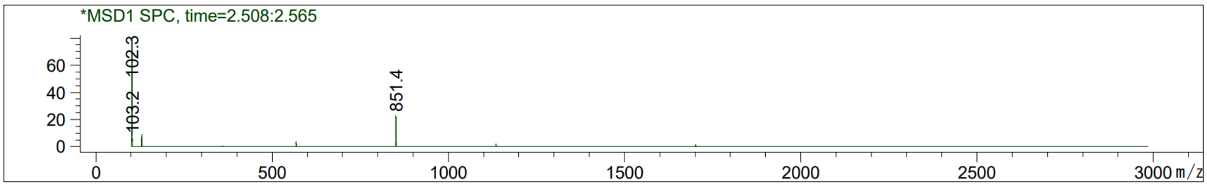

<u>HRMS</u>: Predicted M/z = 1702.54711; Experimental M/z = 1702.54660; Accuracy = 0.30 ppm

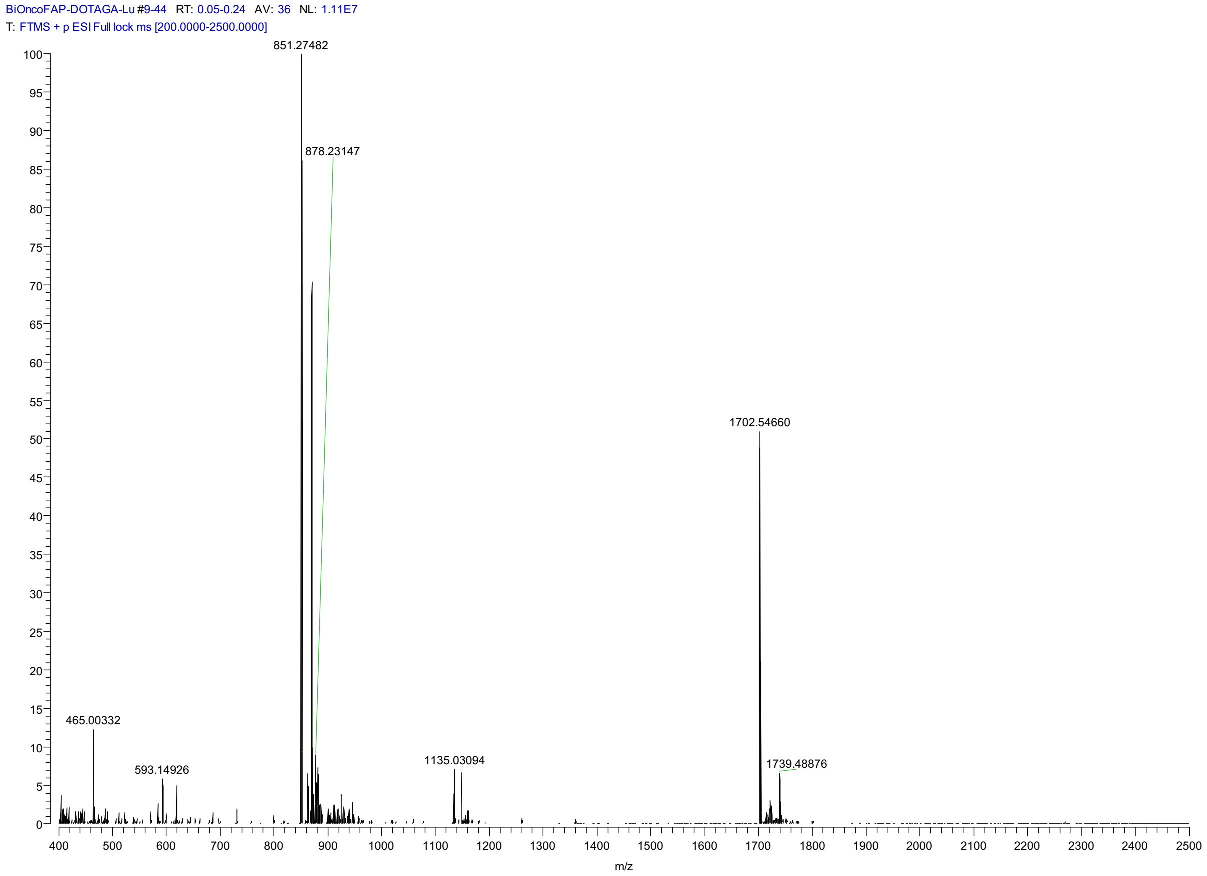

**Synthesis of BiOncoFAP-Asp-Lys-Asp-Cys (15)**

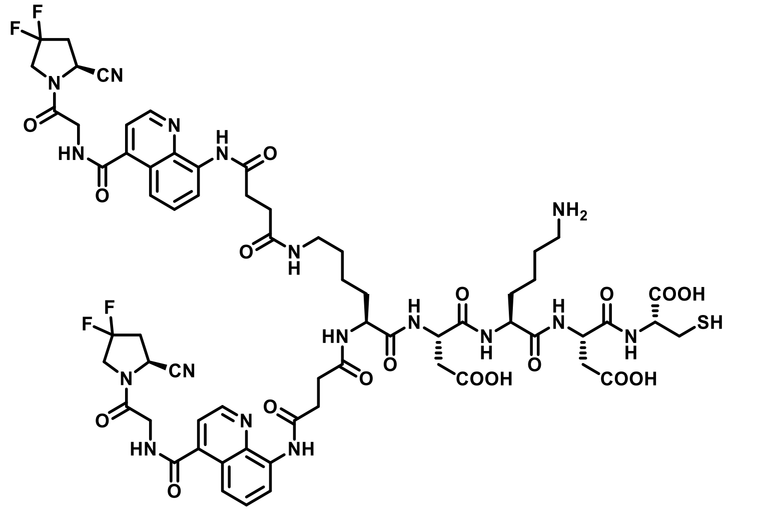

To a solid-phase synthesis syringe, H-Cys(Trt)-2-CT-polystyrene resin (900 mg) was added and then swollen with DMF for 15 min. Fmoc-L-Asp(OtBu)-OH (444 mg, 1.08 mmol, 2 eq.), HATU (411 mg, 1.08 mmol, 2 eq.) and DIPEA (377 µL, 2.16 mmol, 4 eq.) were sequentially added to the resin. The mixture was allowed to react for 2 h, then treated with a 20% solution of Piperidine in DMF (10 mL) for the Fmoc-removal and washed several times with DMF. The resin was then treated with a solution of Fmoc-L-Lys(Boc)-OH (506 mg, 1.08 mmol, 2 eq.), HATU (411 mg, 1.08 mmol, 2 eq.) and DIPEA (377 µL, 2.16 mmol, 4 eq.) in DMF (10 mL) for 2 h, following Fmoc-removal with a 20% solution of Piperidine in DMF (10 mL). After washing with DMF, a solution of Fmoc-L-Asp(OtBu)-OH (444 mg, 1.08 mmol, 2 eq.), HATU (411 mg, 1.08 mmol, 2 eq.) and DIPEA (377 µL, 2.16 mmol, 4 eq.) in DMF (10 mL) was added to the resin. After 1 h, the resin was washed and treated with a 20% solution of Piperidine in DMF (10 mL). Subsequently, Fmoc-L-Lys(Fmoc)-OH (647 mg, 1.08 mmol, 2 eq.), HATU (411 mg, 1.08 mmol, 2 eq.) and DIPEA (377 µL, 2.16 mmol, 4 eq.) and DMF (10 mL) were added to the resin and the mixture was allowed to react for 2 h, following Fmoc-removal with 20% solution of Piperidine in DMF (10 mL). Lastly, the resin was treated with a solution of OncoFAP-COOH (992 mg, 2.16 mmol, 4 eq.), HATU (822 mg, 2.16 mmol, 4 eq.) and DIPEA (754 µL, 4.32 mmol, 8 eq.) in DMF (15 mL) for 2 h. The peptide was then cleaved from the resin using 15 mL of a solution of TFA/Triisopropylsilane/Thioanisol/water in DCM (30 : 5 : 2.5 : 2.5 : 60) for 1 h. The residue was concentrated under *vacuum*, resuspended in cold diethyl ether and centrifugated. The supernatant was discarded, and the pellet was dissolved in DMF and purified via RP-HPLC using a gradient of Water/ACN + 0.1% TFA in 7 min. The desired fractions were collected and lyophilized to afford a white solid. (136 mg, 17%)

MS(ES+) m/z 1490.5 $(M+H)^+$

##### Synthesis of BiOncoFAP-Fluorescein (8)

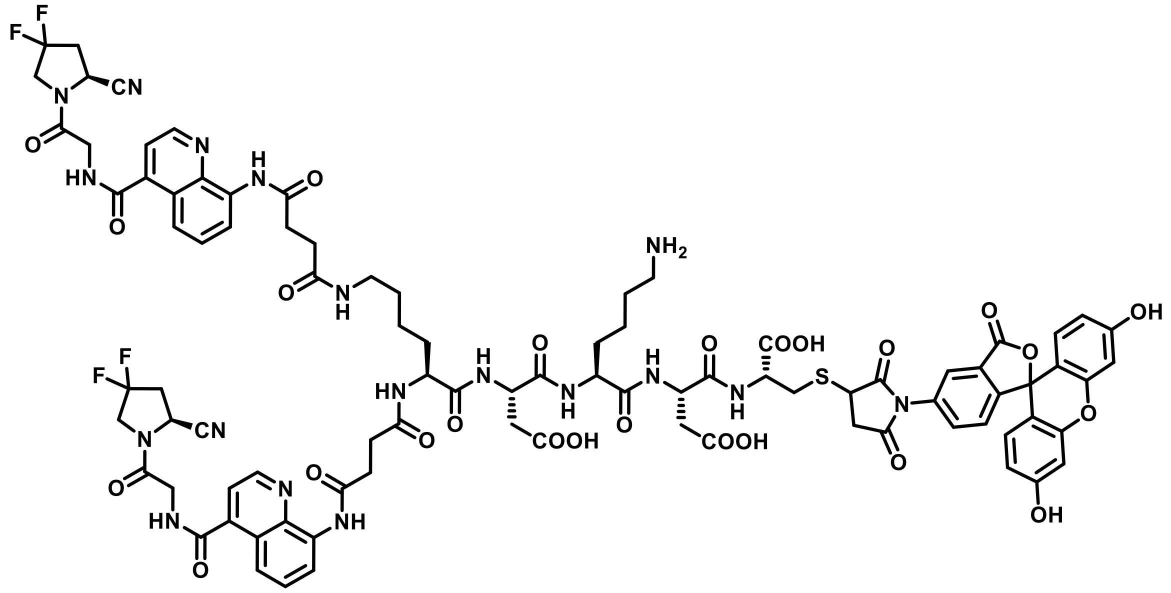

BiOncoFAP-Asp-Lys-Asp-Cys (1.00 mg, 0.59 µmol, 1.0 eq) is dissolved in PBS pH 7.4 (840 µL). Maleimido-Fluorescein (0.76 mg, 1.77 µmol, 3.0 eq) is added as dry DMF solution (160 µL). The reaction is stirred for 3 h. The crude material is purified by RP-HPLC (Water 0.1% TFA/ACN 0.1%TFA 95:5 to 2:8 in 20 min) and lyophilized, to obtain a yellow solid. (1.0 mg, 88%)

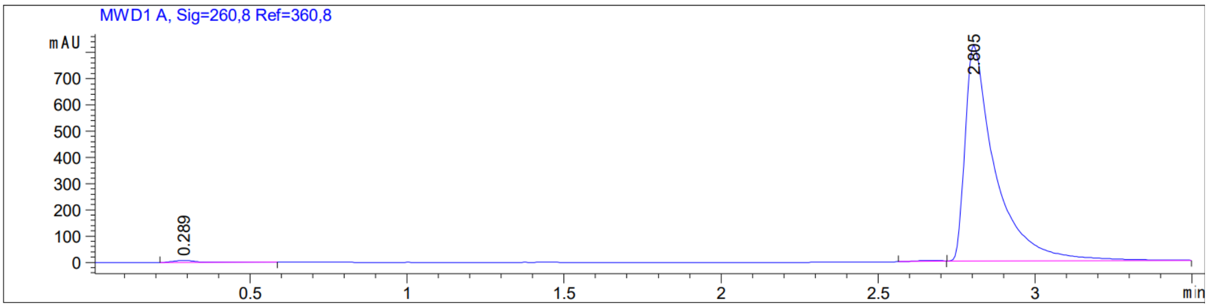

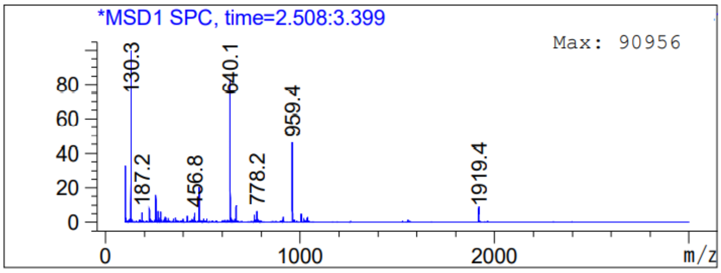

#### Synthesis of BiOncoFAP-Alexa488 (10)

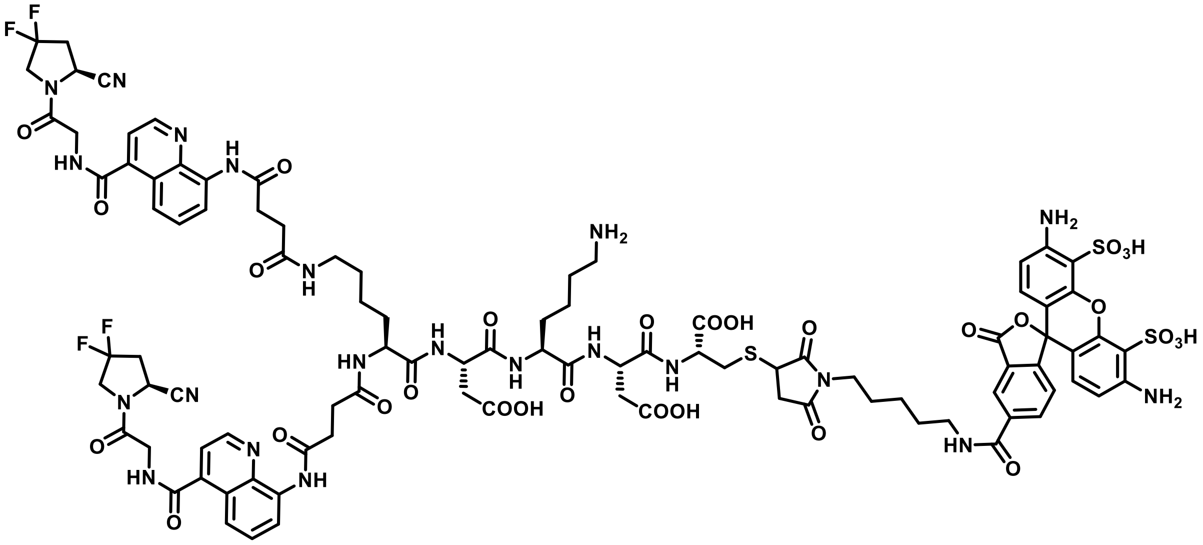

BiOncoFAP-Asp-Lys-Asp-Cys (1.0 mg, 0.59 µmol, 1.0 eq) is dissolved in PBS pH 7.4 (300 µL). Alexa FluorTM 15 488 C5 Maleimide (200 µg, 0.29 µmol, 0.5 eq) is added as dry DMSO solution (200 µL). The reaction is stirred for 3 h. The crude material is purified by RP-HPLC (Water 0.1% TFA/ACN 0.1%TFA 95:5 to 2:8 in 20 min) and lyophilized, to obtain an orange solid. (1.1 mg, 88%)

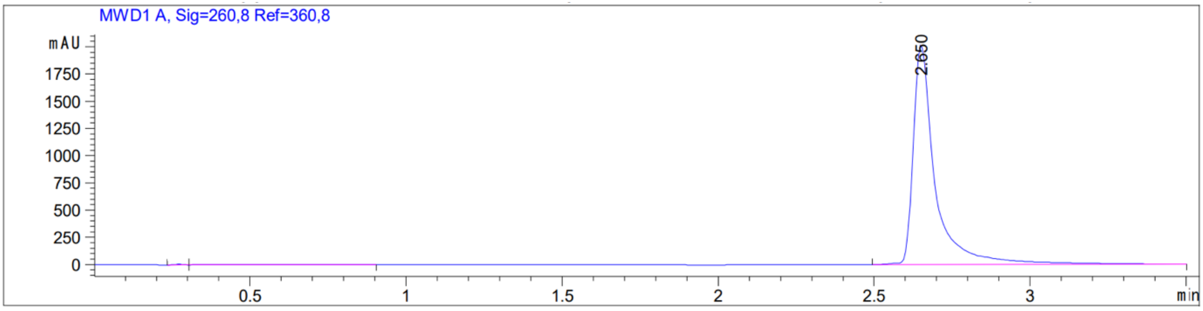

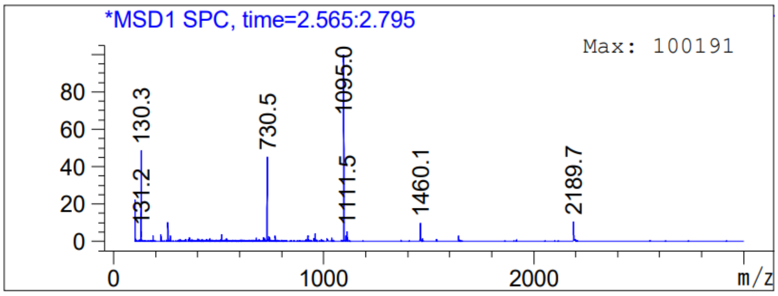

#### Synthesis of BiOncoFAP-IRDye750 (12)

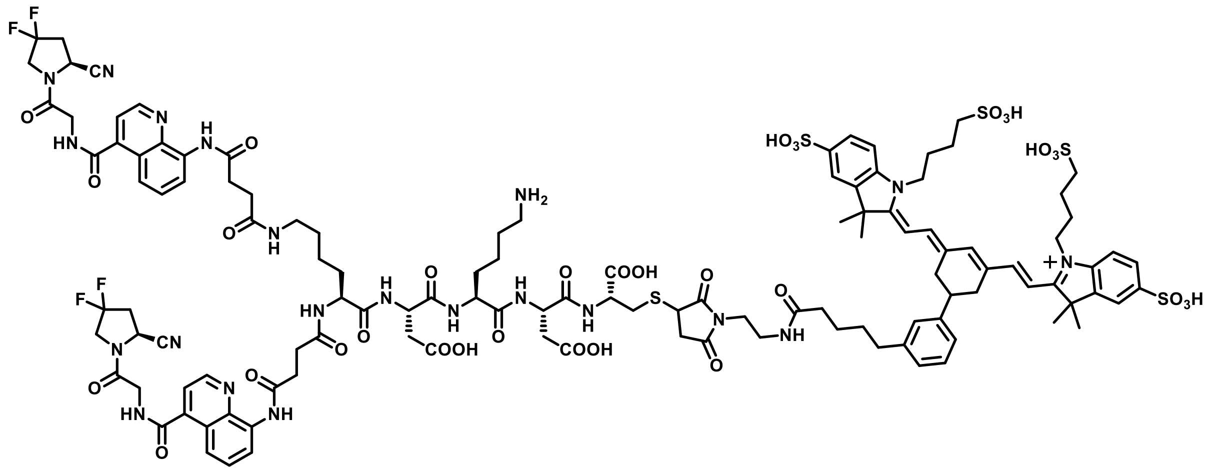

To a solution of BiOncoFAP-Asp-Lys-Asp-Cys (204 µg, 0.14 µmol, 1 eq.) in PBS pH=7.4 (200 µL) was added a solution of IRDye750 maleimide (150 µg, 0.12 µmol, 0.9 eq.) in DMSO (150 µL). The mixture was stirred at room temperature for 3 h and purified via RP-HPLC using a gradient of 90:10 to 50:50 water/ACN + 0.1% TFA in 7 min). The desired fractions were collected and lyophilized to afford a blue solid collect the desired fractions and lyophilize to afford a green solid. (0.2 mg, 54%)

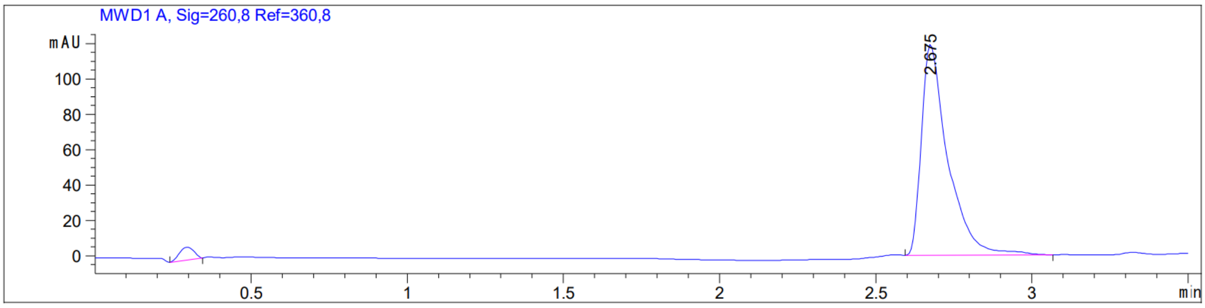

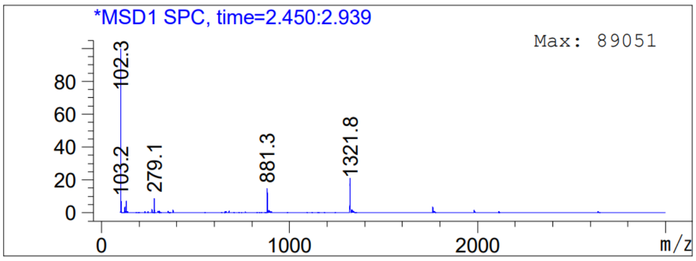

**Synthesis of tert-butyl (8-aminoquinoline-4-carbonyl)glycinate (16)**

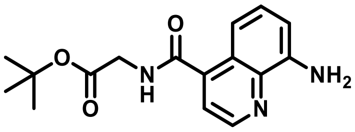

8-aminoquinoline-4-carboxylic acid (200 mg, 1.06 mmol, 1 eq.), HATU (401 mg, 1.06 mmol, 1 eq.) and Gly-OtBu*HCl (214 mg, 1.28 mmol, 1.2 eq.) were suspended in a 4:1 mixture of DCM/DMF (4 mL) and cooled to 0°C. DIPEA (0.74 mL, 4.25 mmol, 4 eq.) was added dropwise to the reaction mixture and stirred at room temperature for 30 min. The solution was transferred to a separatory funnel, diluted with DCM and washed with a saturated aqueous solution of $NaHCO_3$ and brine. The organic phase was dried over anhydrous $Na_2SO_4$, filtered and evaporated under reduced pressure to afford a brown oil. The crude was purified via flash column chromatography (100% DCM to 9:1 DCM/MeOH) to afford a yellow foam. (259 mg, 79%).

MS (ESI+), m/z 302.2

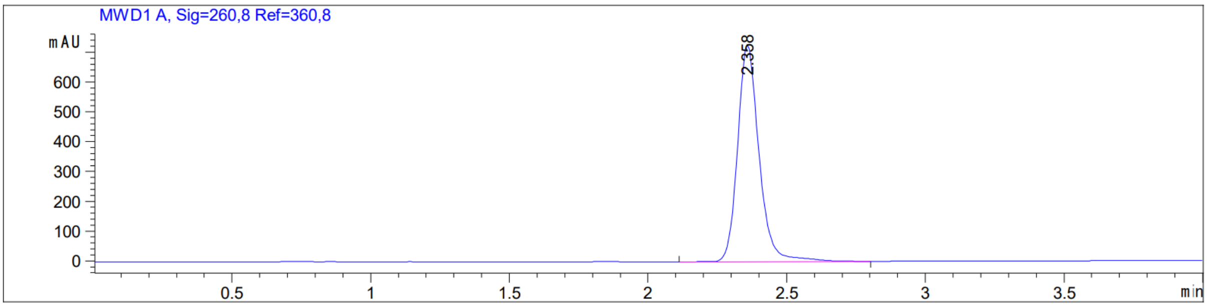

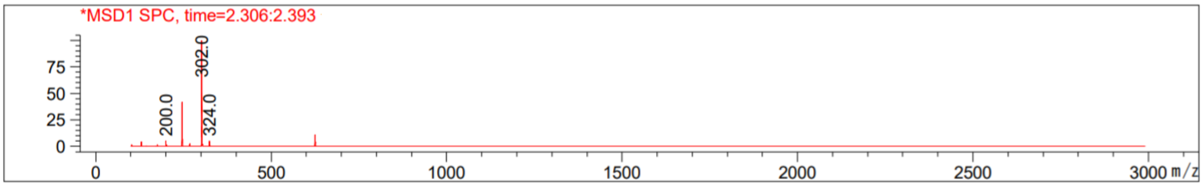

##### Synthesis of negOncoFAP-COOH (17)

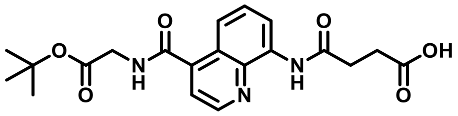

Compound **16** (259 mg, 0.86 mmol, 1 eq.), succinic anhydride (258 mg, 2.58 mmol, 3 eq.) and DMAP (52 mg, 0.43 mmol, 0.5 eq.) were dissolved in dry THF (5 mL). The mixture was heated at 55°C for 1h then cooled to room temperature, diluted with EtOAc and transferred to a separatory funnel. The mixture was washed with brine and the organic phase was dried over anhydrous $Na_2SO_4$, filtered and evaporated under reduced pressure to afford a yellow solid. The crude was purified via flash column chromatography (100% DCM to 8:2 DCM/MeOH) to afford a white solid. (252 mg, 73%)

MS (ESI+), m/z 402.2

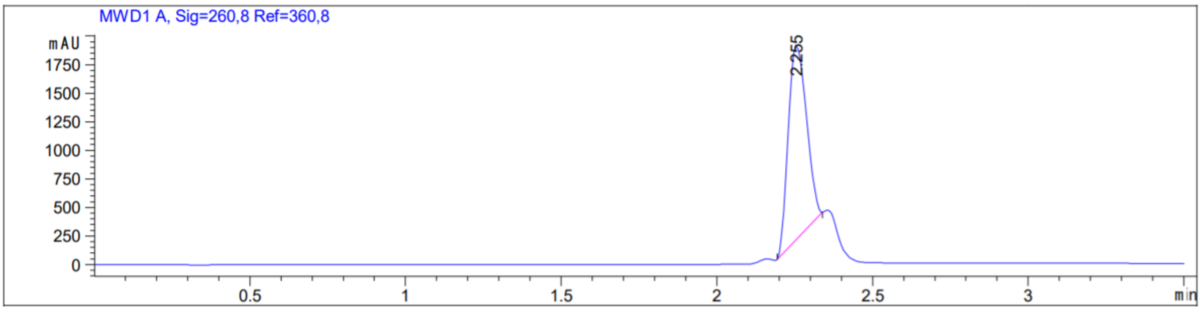

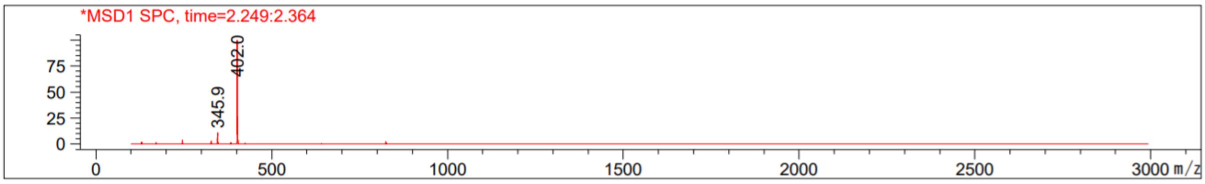

#### Synthesis of negOncoFAP-Asp-Lys-Asp-Cys (18)

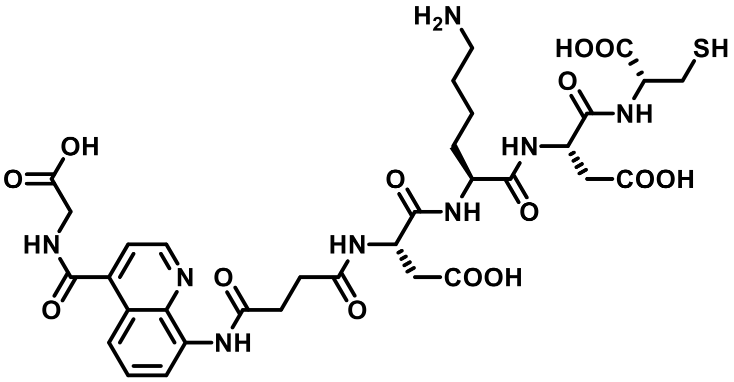

To a solid-phase synthesis syringe, H-Cys(Trt)-2-CT-polystyrene resin (150 mg) was added and then swollen with DMF for 15 min. Fmoc-L-Asp(OtBu)-OH (72 mg, 0.18 mmol, 2 eq.), HATU (67 mg, 0.18 mmol, 2 eq.) and DIPEA (46 µL, 0.36 mmol, 4 eq.) were sequentially added to the resin. The mixture was allowed to react for 2 h, then treated with a 20% solution of Piperidine in DMF (3 mL) for the Fmoc-removal and washed several times with DMF. The resin was then treated with a solution of Fmoc-L-Lys(Boc)-OH (82 mg, 0.18 mmol, 2 eq.), HATU (67 mg, 0.18 mmol, 2 eq.) and DIPEA (46 µL, 0.36 mmol, 4 eq.) in DMF (3 mL) for 2 h, following Fmoc-removal with a 20% solution of Piperidine in DMF (3 mL). After washing with DMF, a solution of Fmoc-L-Asp(OtBu)-OH (72 mg, 0.18 mmol, 2 eq.), HATU (67 mg, 0.18 mmol, 2 eq.) and DIPEA (46 µL, 0.36 mmol, 4 eq.) in DMF (3 mL) was added to the resin. After 1 h, the resin was washed and treated with a 20% solution of Piperidine in DMF (3 mL). Lastly, the resin was treated with a solution of negOncoFAP-COOH (72 mg, 0.18 mmol, 2 eq.), HATU (67 mg, 0.18 mmol, 2 eq.) and DIPEA (46 µL, 0.36 mmol, 4 eq.) in DMF (5 mL) for 2 h. The peptide was then cleaved from the resin using 5 mL of a solution of TFA/Triisopropylsilane/Thioanisol/water in DCM (30 : 5 : 2.5 : 2.5 : 60) for 1 h. The residue was concentrated under *vacuum*, resuspended in cold diethyl ether and centrifugated. The supernatant was discarded, and the pellet was dissolved in DMF and purified via RP-HPLC using a gradient of Water/ACN + 0.1% TFA in 7 min. The desired fractions were collected and lyophilized to afford a white solid. (18 mg, 25%).

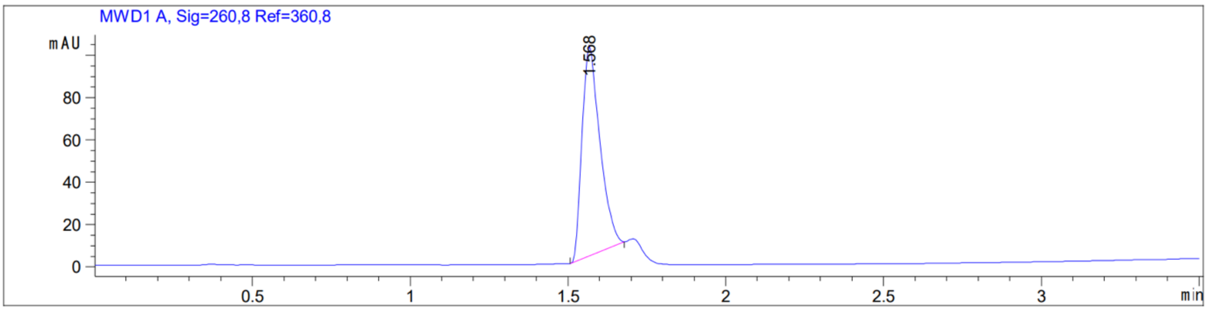

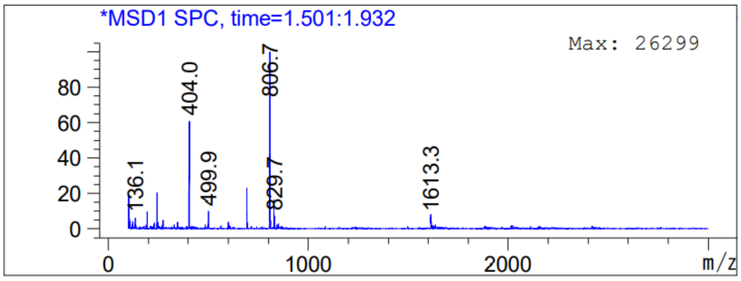
*MSD1 SPC, time=1.501:1.932
Max: 26299
80
60
40
20
0
0
1000
2000
m/z
136.1
404.0
499.9
806.7
829.7
1613.3

#### Synthesis of negOncoFAP-Alexa488 (19)

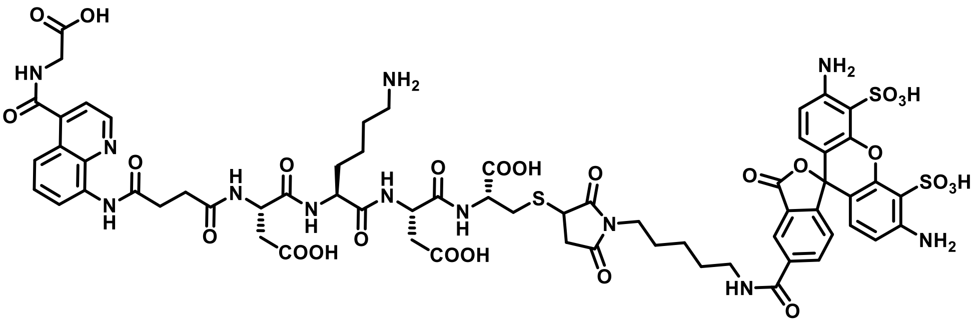

negOncoFAP-Asp-Lys-Asp-Cys (compound **18**, 1.0 mg, 0.6 µmol, 1.0 eq) is dissolved in PBS pH 7.4 (300 µL). Alexa FluorTM 15 488 C5 Maleimide (200 µg, 0.3 µmol, 0.5 eq) is added as dry DMSO solution (200 µL). The reaction is stirred for 3 h. The crude material is purified by RP-HPLC (Water 0.1% TFA/ACN 0.1%TFA 95:5 to 2:8 in 20 min) and lyophilized, to obtain an orange solid. (0.9 mg, 72%)

MS(ES+), m/z 1505.2

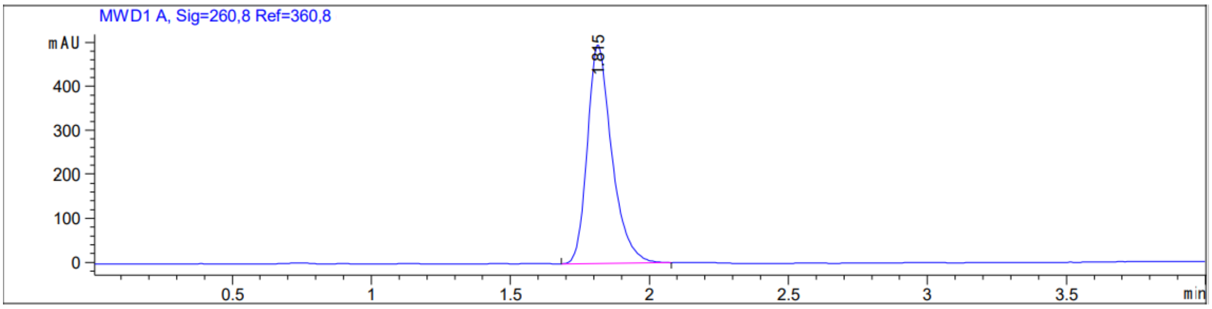

**Synthesis of negBiOncoFAP-Asp-Lys-Asp-Cys (20)**

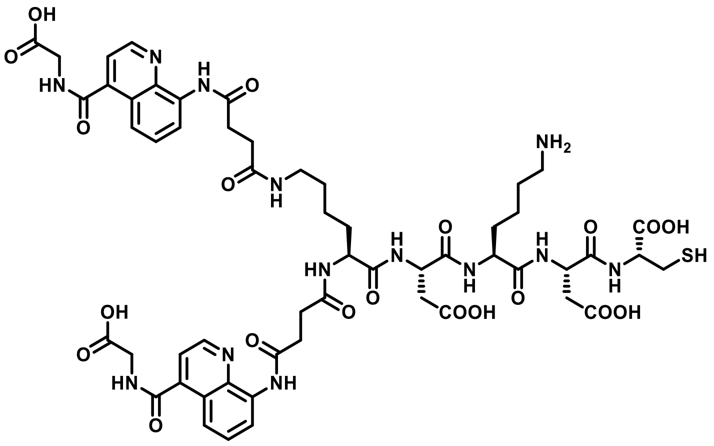

To a solid-phase synthesis syringe, H-Cys(Trt)-2-CT-polystyrene resin (250 mg) was added and then swollen with DMF for 15 min. Fmoc-L-Asp(OtBu)-OH (123 mg, 0.3 mmol, 2 eq.), HATU (114 mg, 0.3 mmol, 2 eq.) and DIPEA (105 µL, 0.6 mmol, 4 eq.) were sequentially added to the resin. The mixture was allowed to react for 2 h, then treated with a 20% solution of Piperidine in DMF (3 mL) for the Fmoc-removal and washed several times with DMF. The resin was then treated with a solution of Fmoc-L-Lys(Boc)-OH (140 mg, 0.3 mmol, 2 eq.), HATU (114 mg, 0.3 mmol, 2 eq.) and DIPEA (105 µL, 0.6 mmol, 4 eq.) in DMF (3 mL) for 2 h, following Fmoc-removal with a 20% solution of Piperidine in DMF (3 mL). After washing with DMF, a solution of Fmoc-L-Asp(OtBu)-OH (123 mg, 0.3 mmol, 2 eq.), HATU (114 mg, 0.3 mmol, 2 eq.) and DIPEA (105 µL, 0.6 mmol, 4 eq.) in DMF (3 mL) was added to the resin. After 1 h, the resin was washed and treated with a 20% solution of Piperidine in DMF (3 mL). Subsequently, Fmoc-L-Lys(Fmoc)-OH (180 mg, 0.3 mmol, 2 eq.), HATU (114 mg, 0.3 mmol, 2 eq.) and DIPEA (105 µL, 0.6 mmol, 4 eq.) and DMF (3 mL) were added to the resin and the mixture was allowed to react for 2 h, following Fmoc-removal with 20% solution of Piperidine in DMF (3 mL). Lastly, the resin was treated with a solution of negOncoFAP-COOH (240 mg, 0.6 mmol, 4 eq.), HATU (228 mg, 0.6 mmol, 4 eq.) and DIPEA (155 µL, 1.20 mmol, 8 eq.) in DMF (15 mL) for 2 h. The peptide was then cleaved from the resin using 5 mL of a solution of TFA/Triisopropylsilane/Thioanisol/water in DCM (30 : 5 : 2.5 : 2.5 : 60) for 1 h. The residue was concentrated under vacuo, resuspended in cold diethyl ether and centrifugated. The supernatant was discarded, and the pellet was dissolved in DMF and purified via RP-HPLC using a gradient of water/ACN + 0.1% TFA in 7 min. The desired fractions were collected and lyophilized to afford a white solid. (24 mg, 13%)

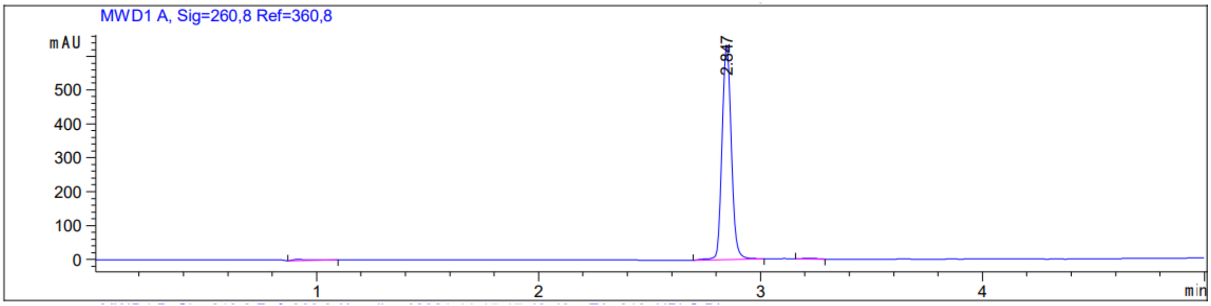
MWD1 A, Sig=260,8 Ref=360,8
mAU
500
400
300
200
100
0
2.847
1
2
3
4
min

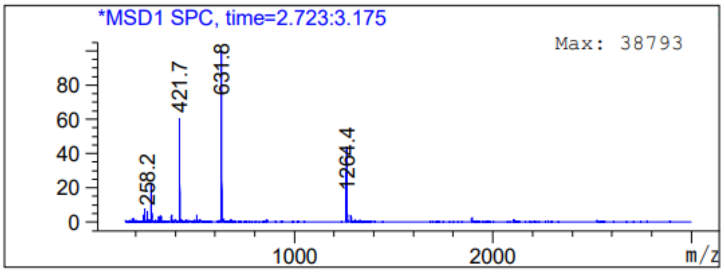
*MSD1 SPC, time=2.723:3.175
Max: 38793
80
60
40
20
0
258.2
421.7
631.8
1264.4
1000
2000
m/z

#### Synthesis of negBiOncoFAP-Alexa488 (21)

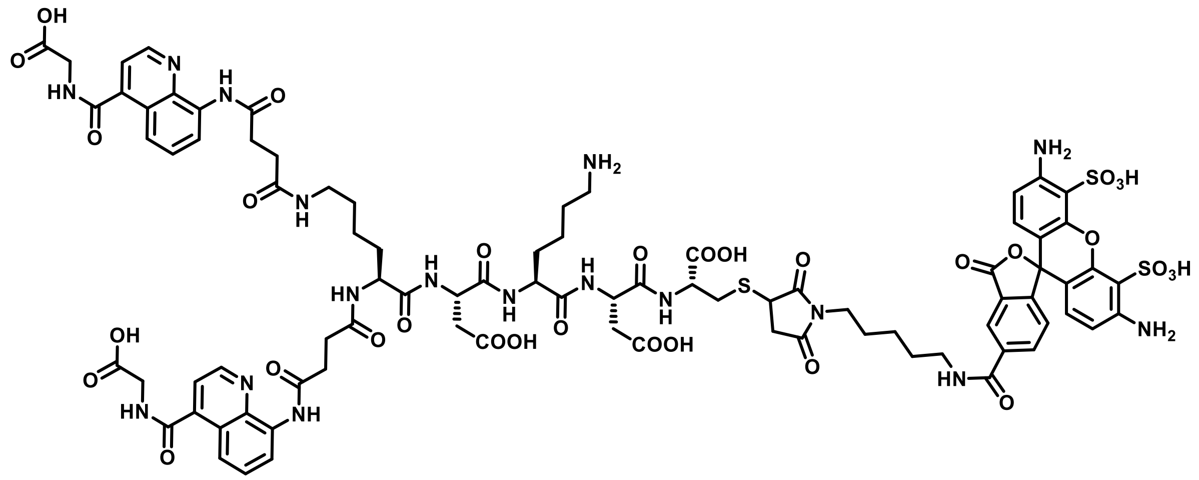

negBiOncoFAP-Asp-Lys-Asp-Cys (compound **22**) (1 mg, 0.59 µmol, 1.0 eq) is dissolved in PBS pH 7.4 (300 µL). Alexa FluorTM 15 488 C5 Maleimide (200 µg, 0.29 µmol, 0.5 eq) is added as dry DMSO solution (200 µL). The reaction is stirred for 3 h. The crude material is purified by RP-HPLC (Water 0.1% TFA/ACN 0.1%TFA 9.5:0.5 to 2:8 in 20 min) and lyophilized, to obtain an orange solid. (0.9 mg, 59%)

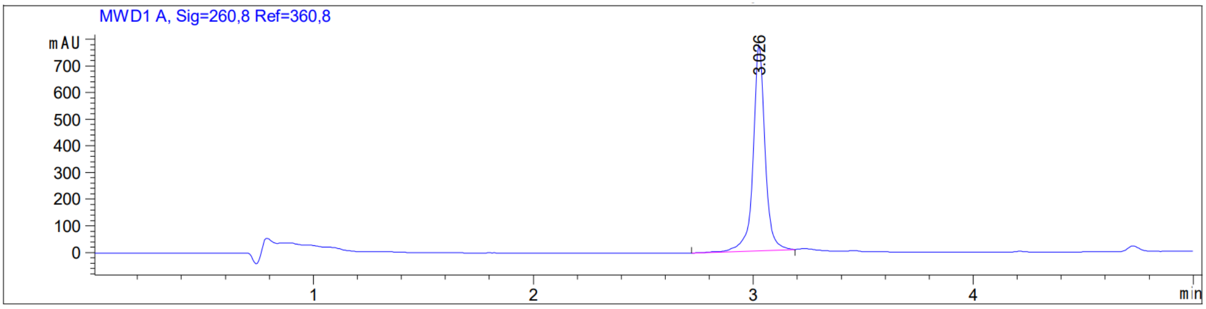

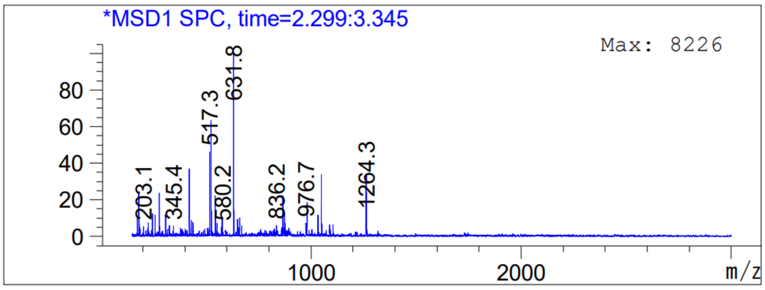

#### Quality control of radiosynthesis – Radio-HPLC

Reversed-phase Radio-HPLC were performed on a Merck-Hitachi D-7000 Series equipped with Raytest GABI-Star radio detector, using a Synergi 4 µm Polar-RP 80 Å, 150 x 4.6 mm column at a flow rate of 1 ml min-1 with linear gradients of solvents Millipore water and CAN.

#### Quality control of radiosynthesis - Coelution Experiments of Ligand–Protein Complexes.

PD-10 columns were pre-equilibrated with running buffer (50 mM Tris, 100 mM NaCl, pH = 7.4). 150 µL of a solution containing hFAP (2 µM) or hCAIX (irrelevant protein, 2 µM) was pre-incubated with 2 µL of $^{177}$Lu-OncoFAP or $^{177}$Lu-BiOncoFAP stock solution (50 µM, 5 MBq). The final solution was loaded on the column and flushed with running buffer. Fractions of the flow-through (200 µL) were collected in test tubes and the radioactivity measured with a Packard Cobra γ-counter. As negative control, 2 µL of $^{177}$Lu-OncoFAP or $^{177}$Lu-BiOncoFAP stock solution (50 µM, 5 MBq) were diluted in 150 µL of running buffer (50 mM Tris, 100 mM NaCl, pH = 7.4), without proteins. The final solution was loaded on the column and flushed with running buffer. Fractions of the flow-through (200 µL) were collected in test tubes and the radioactivity measured with a Packard Cobra γ-counter. Results of the co-elution experiments performed with $^{177}$Lu-OncoFAP and $^{177}$Lu-BiOncOFAP on hFAP, hCAIX (irrelevant protein) and without protein are shown in Figure S1.

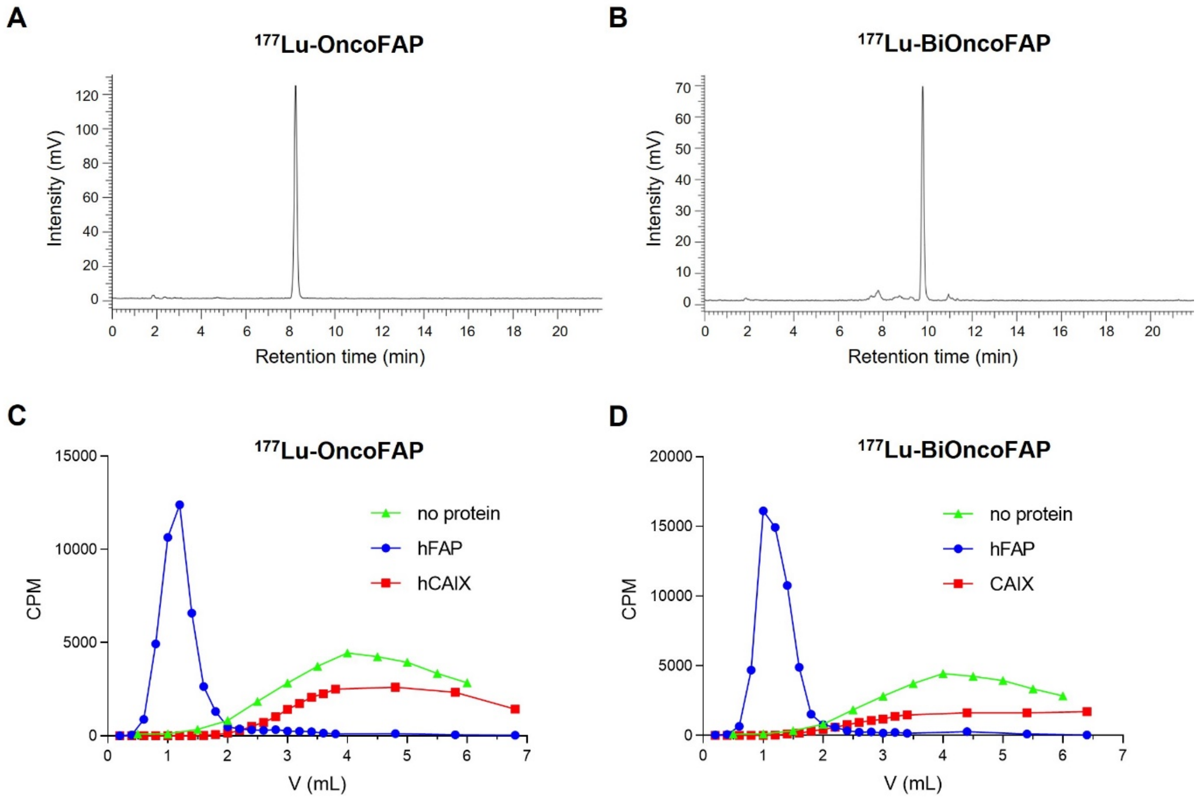

**Figure S1.** Quality control of radiosynthesis. Radio-HPLC of $^{177}$Lu-OncoFAP (*A*) and $^{177}$Lu-BiOncoFAP (*B*); PD-10 experiments of $^{177}$Lu-OncoFAP (*C*) and $^{177}$Lu-BiOncoFAP (D) incubated with hFAP, using no protein or irrelevant protein (hCAIX) as a negative control.

#### *In vitro* tests and assays

##### Affinity Measurement to non-target proteins by Fluorescence Polarization.

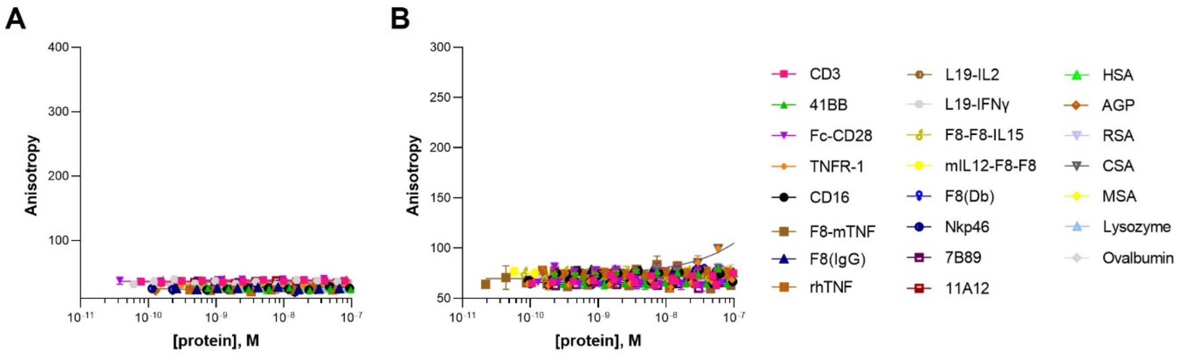

**Figure S2.** Fluorescence polarization experiments of OncoFAP-Fluorescein (*A*) and BiOncoFAP-Fluorescein (*B*) towards a panel of non-target proteins, including tumour-associated antigens, abundant proteins and irrelevant proteins.

##### Stability in human and mouse blood serum.

36 µL of serum (either mouse "Sigma Aldrich", human "Sigma Aldrich") were preincubated at 37 °C for 5 minutes. 4 µL of a 1 mM DMSO solution of the analytes was added at the final concentration of 50 µM to start the kinetic. The assay was blocked by deproteinization with 300 µL of ACN at 0, 24, 48, 72 and 120 hours after the addition of the compound. Deproteinized samples were centrifugated at 14000 g for 10 minutes. 200 µL of supernatant was dried under vacuum at 37 °C and carefully resuspended in 30 µL of an aqueous solution containing 10% ACN and 0.1 % HCOOH. Samples were analyzed via LC-MS. Chromatographic separation was carried out on an Agilent 1200 Series LC System using as column an InfinityLab Poroshell 120 EC-C18, (4.6 x 56 mm) at a flow rate of 0.8 mL/min with linear gradients of solvents A and B (A = Millipore water with 0.1% formic acid, B = ACN with 0.1% formic acid) from 40% to 100% of B in 3 minutes. Eluents were analyzed in full mass scan in positive ion mode with an Agilent 6100 Series Single Quadrupole MS 5 System.

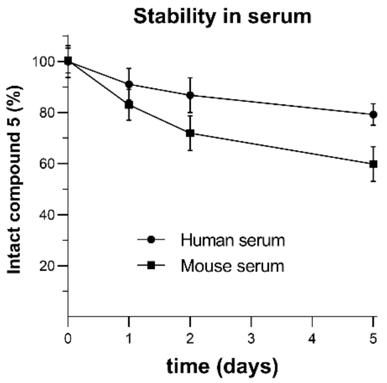

**Figure S3.** *In vitro* stability of [$^{nat}$Lu]Lu-BiOncoFAP-DOTAGA (**5**) in human and mouse serum at 37°C, 50 µM over time.

**Determination of Log$D_{7.4}$ values of $^{177}$Lu-OncoFAP and $^{177}$Lu-BioncoFAP.**

The lipophilicity of $^{177}$Lu-OncoFAP and $^{177}$Lu-BiOncoFAP was determined as follows. 100 µL aliquotes of the radioligand (~ 1 MBq) in PBS buffer were added to PBS buffer (500 µL, pH 7.4) and 1-octanol (600 µL). The two-layer mixture were vigorously shaken for 10 minutes on a vortex mixer and then centrifuged at 700 rpm for 5 min to facilitate the separation. 100 µL aliquotes of both layers were measured in a Packard Cobra Gamma Counter and the partition coefficient was determined by dividing cpm (octanol) by cpm (PBS) and indicated as Log$D_{7.4}$.

Log$D_{7.4}$ ($^{177}$Lu-OncoFAP): -4.02

Log$D_{7.4}$ ($^{177}$Lu-BiOncoFAP): -3.60

### Animal studies

##### *In vivo* Tumour and Organ Penetration Analysis of OncoFAP and BiOncoFAP.

SK-RC-52.hFAP and SK-RC-52.wt xenografted tumours were implanted respectively into the right and left flank of female athymic Balb/c AnNRj-Foxn1 mice (6-8 weeks of age) as described above, and allowed to grow to an average volume of 200 $mm^3$.

OncoFAP-IRDye750 (compound **11**) and BiOncoFAP-IRDye750 (compound **12**) were administered intravenously at a dose of 250 nmol/kg.

Fluorescence images were aquired at different time points (10 min, 1 h, 2 h, 3 h, 4.5 h, 6 h) on an IVIS Spectrum imaging system (Xenogen, exposure 1s, binning factor 8, excitation at 745 nm, emission filter at 800 nm, f number 2, field of view 13.1). Six hours after administration, mice were euthanized by $CO_2$ asphyxiation. Tumours, organs and blood were collected, and fluorescence images acquired as described above.

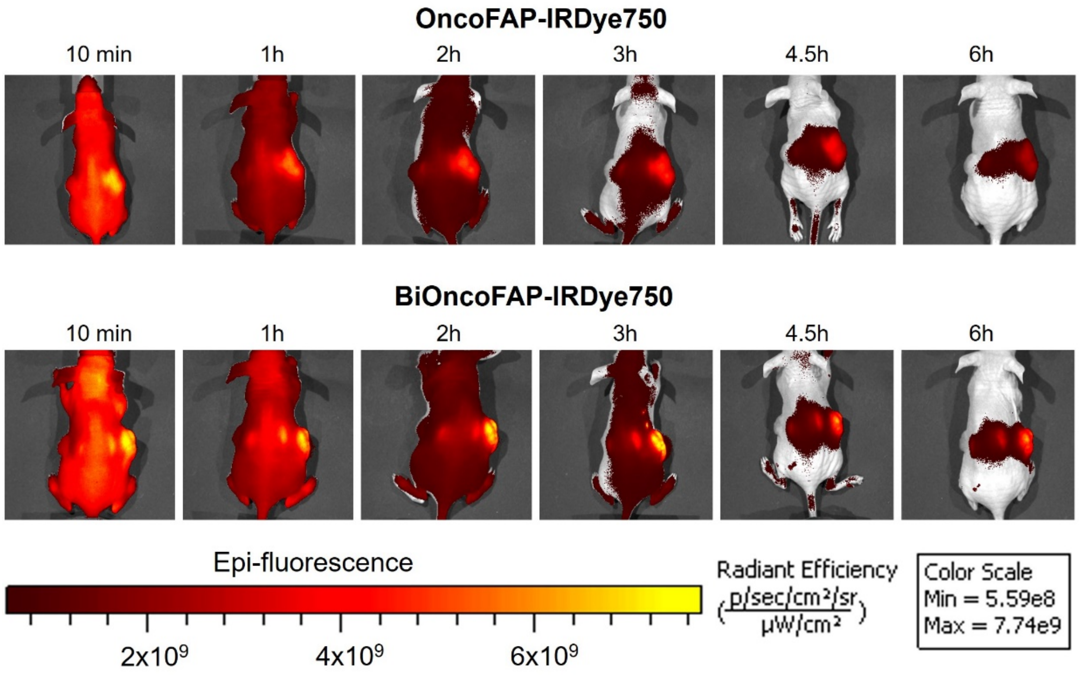

**Figure S4.** Near-infrared fluorescence imaging evaluation of the targeting performance of OncoFAP-IRDye750 and BiOncoFAP-IRDye in mice bearing SK-RC-52.hFAP tumours (right flank) and SK-RC-52.wt tumours (left flank). Images were collected at different time points (10 min, 1 h, 2 h, 3 h, 4.5 h, 6 h) after the intravenous injection (250 nmol/kg).

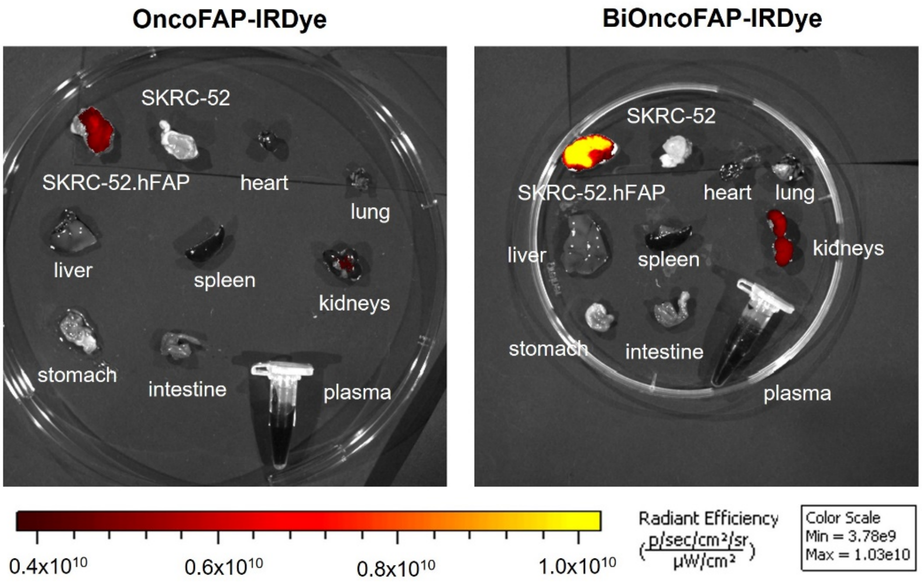

**Figure S5.** Near-infrared fluorescence imaging evaluation of the targeting performance of OncoFAP-IRDye750 and BiOncoFAP-IRDye in mice bearing SK-RC-52.hFAP tumours (right flank) and SK-RC-52.wt tumours (left flank). Mice were euthanized 6 h after systemic administration (250 nmol/kg) and images were collected.
